# Identifying genetic determinants of complex phenotypes from whole genome sequence data

**DOI:** 10.1101/181222

**Authors:** George S. Long, Mohammed Hussen, Jonathan Dench, Stéphane Aris-Brosou

## Abstract

A critical goal in biology is to relate the phenotype to the genotype, that is, to find the genetic determinants of various traits. However, while simple monofactorial determinants are relatively easy to identify, the underpinnings of complex phenotypes are harder to predict. While traditional approaches rely on genome-wide association studies based on Single Nucleotide Polymorphism data, the ability of machine learning algorithms to find these determinants in whole proteome data is still not well known. To better understand the applicability of machine learning in this case, we implemented two such algorithms, adaptive boosting (AB) and repeated random forest (RRF), and developed a chunking layer that facilitates the analysis of whole proteome data. We first assessed the performance of these algorithms and tuned them on an influenza data set, for which the determinants of three complex phenotypes (infectivity, transmissibility, and pathogenicity) are known based on experimental evidence. This allowed us to show that chunking improves runtimes by an order of magnitude. Based on simulations, we showed that chunking also increases sensitivity of the predictions, reaching 100% with as few as 20 sequences in a small proteome as in the influenza case (5k sites), but may require at least 30 sequences to reach 90% on larger alignments (500k sites). While RRF has less specificity than RF, it was never < 50%, and RRF sensitivity was significantly higher at smaller chunk sizes. We then used these algorithms to predict the determinants of three types of drug resistance (to Ciprofloxacin, Ceftazidime, and Gentamicin) in a bacterium, *Pseudomonas aeruginosa*. While both algorithms performed well in the case of the influenza data, results were more nuanced in the bacterial case, with RRF making more sensible predictions, with smaller errors rates, than AB. Altogether, we demonstrated that ML algorithms can be used to identify genetic determinants in small proteomes (viruses), even when trained on small numbers of individuals. We further showed that our RRF algorithm may deserve more scrutiny, which should be facilitated by the decreasing costs of both sequencing and phenotyping of large cohorts of individuals.

## Background

An overarching goal in biology is to predict an individual’s phenotype from its genotype, in a given environment [1], or from a given genetic makeup [2]. An obvious first step in this direction is to find the genetic determinants of each phenotype of interest – which is what genome-wide association studies (GWAS’s) have endeavored to achieve over the past ten years [3]. At their foundation, GWAS’s rely on the analysis of millions of variants in the genome, without any prior knowledge of their involvement with a particular phenotype, over a sample of unrelated individuals: as such, GWAS’s are often qualified of performing an “unbiased scan of the genome” [4]. While GWAS’s have some limitations that may be shared by alternative approaches (they assume that common diseases are caused by common variant [4], which may result in failing to explain most of the phenotypic variance [5]), their main issue may lie in their reliance on Single Nucleotide Polymorphism (SNP) data, that only offer a partial snapshot of the genome, and that may not even contain the causative agents of the phenotype under study. The immediate alternative to using SNP data would be resort to high throughput sequencing technology, and work directly with entire proteome data [6], which are predicted to replace SNP data [3]. However, changing from focal SNP’s to whole proteome information will radically increase the number of tests to be performed, and might lead to statistical complications in controlling false discovery, as acknowledged at the very outset of GWAS [7]. Recently, it was suggested that machine learning could be used to predict susceptibility to some cancers in humans [8]. Building upon these developments, we here hypothesized that machine learning is able to go beyond predicting genomic inheritance within a cohort of individuals, by predicting the genetic determinants of particular phenotypes using proteome data.

A large number of machine learning approaches exist [9], and are gaining popularity in biology [10, 11]. For instance, in the work just cited above, the authors resorted to neural networks [8]. While some statisticians claim that adaptive boosting (AB) [12] is the “best off-the-shelf classifier in the world” [13, 9], it is probable that no single classifier can be considered perfect in all situations – a situation known as the *no free lunch theorem* [14]. Technically, AB relies on an iterated process where linear decisions are fitted in the space of proteomic features (amino acid positions in a protein alignment). At each iteration, individual observations that were misclassified in the previous iteration are emphasized, so that the algorithm learns from past errors. While each iteration typically leads to a weak classifier that is just a bit better than chance, the final classifier takes advantage of combining these weak classifiers to improve (*boost*) their performance and construct a strong one [12], *i.e*. a classifier with an accuracy that can come close to 99% [15]. While our recent success with AB [16] prompted us to further investigate this algorithm in the context of proteome data, we also aimed at comparing its performance with more popular algorithms, such as random forests [17] (RF). The RF algorithm is essentially based on decision trees [18], combined with a bootstrap procedure (bagging) aimed at increasing the stability of the predictions, and a random subsetting of predictors to decorrelate the bagged trees, that are then averaged to produce the final classifier.

As any supervised learning algorithm, AB and RF are trained on labeled data, *i.e*. data for which the correct assignments are known. In our case, the proteome data coming from an organism for which drug resistance (the label / phenotype) is known. Before training the algorithm, the labeled data are usually split into two subsets, a *train* and a *test* data set, where only the former is used to train the classifier. This trained classifier is then tested on the independent *test* data set, to validate its performances, and can then be used to classify new individuals, which were neither part of the *train* nor of the *test* data sets, as being “drug resistant” or not, based solely on their proteomic information. Here however, our goal is slightly different: we are not interested in classifying individuals, but in finding the most important features (again, these are the mutations at particular sites) that the algorithm is learning from to correctly classify individuals. This is why training will be done on the entire data, rather than splitting them as *train* and *test* data sets. More specifically, during the learning process, features (sites) are ranked by decreasing importance in fitting the final model [19]. The algorithm weighs the influence of each feature based on their relative importance during the creation of the model [20], and thus uses only the most important sites. The end result is that we have not merely a model for predicting phenotype, but a means of identifying a ranked list of the most important features [10] that determine a particular phenotype.

To assess the ability of machine learning algorithms to predict the genetic determinants of particular phenotypes, we implemented two such algorithms, AB and a modified RF, and compared their performance in two microbes, one for which most of these determinants are known (the Influenza A virus), and one for which little is known (the *Pseudomonas aeruginosa* bacterium). In both cases, we retrieved complete proteome sequences of phenotyped individuals. These phenotypes pertained either to their capacity to infect a human host (influenza), or their resistance to particular antibiotics (pseudomonas). Although these two biological systems are very different, both are expected to have a highly complex genetic basis: the molecular determinants of influenza virulence and pathogenesis can span its entire genome [21, 22], and bacterial drug resistance can involve many distinct mechanisms [23]. We took advantage of an influenza database backed by experimental validations [24], and of a recent study of *P. aeruginosa* genomics [25], to train both algorithms. We modified both original algorithms to make them amenable to analyzing whole proteome data sets, and altered the RF algorithm to further stabilize its predictions by introducing a Repeated Random Forest (RRF) algorithm. We then evaluated the performance of these algorithms with respect to either experimental validations (influenza), or both gene annotations and cross-validations (pseudomonas). We discussed the advantages and limitations of our modified algorithms in identifying the genetic determinants of complex phenotypes from whole genome sequence data.

## Results and Discussion

### Determination of the RRF thresholds on the influenza data

The influenza data represent our gold standard here, as we can run our machine learning algorithms on them, and compare the model predictions with experimental evidence (Table 1). Because both algorithms (AB and RRF) essentially return a list of sites by decreasing importance, we can use the influenza data to determine optimal thresholds for predicting genuine genetic determinants. Two thresholds were employed here: the percentile and the consensus thresholds (AB only has the former). This was done by maximizing the number of true positives, *i.e*. sites that (i) sites that are found in at least a certain number of RF runs of the RRF (our “consensus threshold”) – given of course that these sites are backed by experimental evidence in the case of the influenza data (Table 1) – and (i) sites have a high importance or mean Gini index (our “percentile threshold”; Figure S1). By varying both thresholds simultaneously, we determined that using a consensus threshold around 50% and the 90^th^ percentile (top 10% important sites) maximized the number of true positives while minimizing the false positives (Figures S2-S11). Increasing the percentile threshold to 95% dramatically increased false negatives, while decreasing it to the 85th percentile only led to more false positives. Likewise, we found that increasing the consensus threshold only decreased the proportion of false positives, without affecting sensitivity. We henceforth used these two RRF thresholds (50% consensus, 90^th^ percentile).

**Table 1.** Sensitivity of the algorithms on the analysis of the influenza data. This table lists the genes and amino acid positions known to be involved in the three phenotypes studied here, and which one of these were rediscovered by our algorithms. For AB, chunk sizes of 75, 125, and 175 were used to calculate the importance values of each site for adaptive boosting. An importance threshold of 1 was used to determine whether a site was a potential genetic determinant. For RRF, chunk sizes of 80, 125, and 175 were used with a threshold of the 90^th^ percentile and a 60% consensus. Data on experimental validations are from the Influenza Research Database [24]. Genes are ordered by segment size. See Figures 2 and 3 for the specificity of these algorithms.

| Site | AB | RRF | Phenotype | References |
| --- | --- | --- | --- | --- |
| PB2 9 | ✓ | ✓ | Infectivity | [61] |
| PB2 105 | ✓ | ✓ | Pathogenicity/Infectivity | [62] |
| PB2 339 | ✓ | ✓ | Infectivity | [63, 64] |
| PB2 391 | ✓ |  | Transmissibility | [65] |
| PB2 627 | ✓ | ✓ | Infectivity | [66] |
| PB2 667 | ✓ |  | Infectivity | [67] |
| PB1 215 | ✓ | ✓ | Pathogenicity | [68] |
| PB1 375 |  | ✓ | Pathogenicity | [69] |
| PB1 757 | ✓ |  | Infectivity | [70] |
| HA 163 |  | ✓ | Pathogenicity/Infectivity | [71] |
| HA 212 |  | ✓ | Pathogenicity/Infectivity | [72] |
| HA 246 | ✓ | ✓ | Transmissibility | [73] |
| HA 536 |  | ✓ | Infectivity | [74] |
| NP 400 |  |  | Pathogenicity | [75] |
| NA 49 |  | ✓ | Transmissibility | [76] |
| NA 75 |  |  | Transmissibility | [77] |
| M2 31 |  | ✓ | Pathogenicity/Infectivity | [76] |
| NS1 127 |  |  | Pathogenicity | [78] |
| NS1 195 |  |  | Transmissibility/Infectivity | [79] |
| NS1 212 |  | ✓ | Pathogenicity/Infectivity | [80] |

### Chunking improves runtimes, minimally affecting site ranking

As these algorithms can have large computational requirements for whole proteome analyses (AB in particular), we implemented a chunking algorithm (Figure S12). For this, each alignment was subdivided into smaller alignments (*chunks*), on which each ML algorithm was run in a first pass to determine a set of positions of interests (AB: those with importance > 1.5; RRT: those with a Gini index > 0.075). A second pass ran the same ML algorithm on the entire set of positions of interests to determine the final important sites or selected features over the entire data set. The consensus and percentile thresholds described above are then applied to produce the list of genetic determinants (Figure S1).

To determine how chunking affected both the runtimes and the predictions of the machine learning algorithms on the influenza data, 20 chunk sizes were compared, ranging from 75 (or 80 in the case of RRF) to 175 by increments of five. Being much faster than AB, the RRF algorithm was further tested beyond the initial 20 chunks, up until the sequence alignment was analyzed in full. This was repeated for each of the three influenza phenotypes. As expected from its associated increased memory requirements, increasing chunk sizes also increased runtimes exponentially for both AB (Figure 1A) and RRF (Figure 1B). Smaller chunk sizes reduced runtimes by about an order of magnitude (1 log_10_ unit) for both AB and RF. More specifically, an analysis of covariance (ANCOVA) showed that runtimes were similarly affected across all phenotypes in AB (*P* = 0.855). However, while the slopes of these regressions were similar, the analyses for the pathogenicity phenotype ran the slowest (intercept: 3.85; *P* = 1.12 × 10^−5^), followed by transmissibility (intercept: 2.68; *P* = 9.26 × 10^−7^) and infectivity (intercept: 2.35; *P* = 4.61 × 10^−6^). Similar results were observed with RRF, where again all slopes were similar (*P* = 0.973), pathogenicity ran the slowest (intercept: 1.175, *P* < 2.00 × 10^−16^), followed by infectivity (intercept: 1.174, *P* < 2.00 × 10^−16^) and transmissibility (intercept: 1.168, *P* < 2.00 × 10^−16^). As the feature space was the same (the same amino acid alignment), it must be the distribution of the phenotypes (*labels*) that impacted runtimes. However, base runtimes (*i.e*., their intercepts) did not increase due to class imbalance, as the most imbalanced phenotype (transmissibility; Table 5) had an intermediate intercept (Figure 1A). Finally, note that RRF was actually so fast at small chunk sizes that there was a significant overhead associated with our parallelization of the algorithm (Methods), as linear models fitted to only chunk sizes less than 200 were highly significant (*P* < 2.00 × 10^−16^), with negative slopes (Figure 1B).

**Figure 1.**
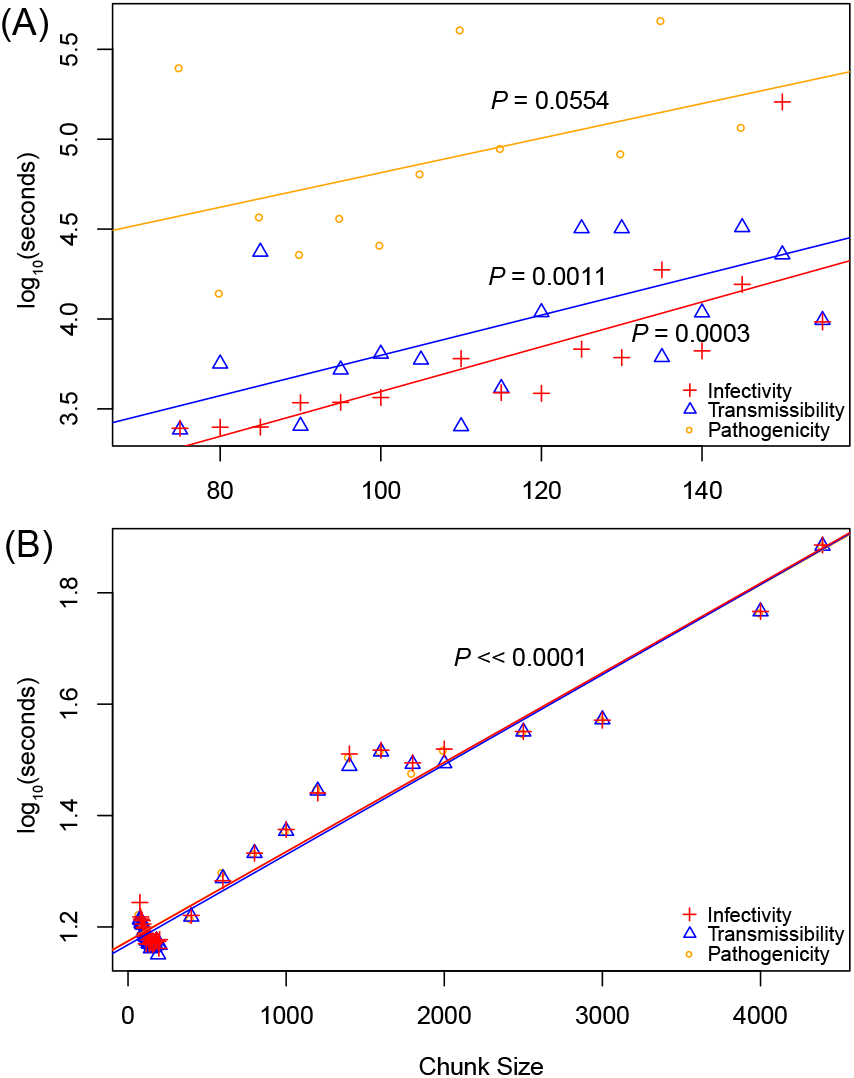
Impact of chunk size on the runtime of the machine learning algorithms for the influenza data. Runtimes for infectivity (red), transmissibility (blue), and pathogenicity (orange) are shown for AB (A) and RRF (B). While each data point is based on a single run, run-to-run variability is taken into account by performing linear regressions (solid lines); their *P*-values are also shown.

Not only were runtimes significantly and similarly reduced by chunking across phenotypes and algorithms, but differences in terms of which amino acids were predicted to be the most important were also similar (Figures 2–3; see Venn diagrams in insets, and distributions of importance values for the largest chunk size). In the case of AB, the six most important sites determining infectivity at the smallest chunk size (75) were among the top fifteen sites at intermediate (125) and largest (175) chunk sizes (Figure 2A). Furthermore, the top site, HA 108, was the most important at all of these chunk sizes, while PB2 627 and 667 were always ranked second or third. The RRF results showed a similar pattern in terms of which sites were the most important (Figure 3). In the case of infectivity, the four top sites (HA 108, HA 536, NA 84 and NA 340) were consistently the most important. After the fourth site, a large drop in importance was observed, suggesting that limited information was available at those sites. But while this drop was also observed for transmissibility at extreme chunk sizes, it was not observed anywhere else, suggesting that the first large drop in ranked importance should not be used to evaluate the relative merit of these predictions. As expected from its repeated nature, the RRF predictions were more stable than those under AB across chunk sizes (compare Venn insets in Figures 3 and 2, respectively).

**Figure 2.**
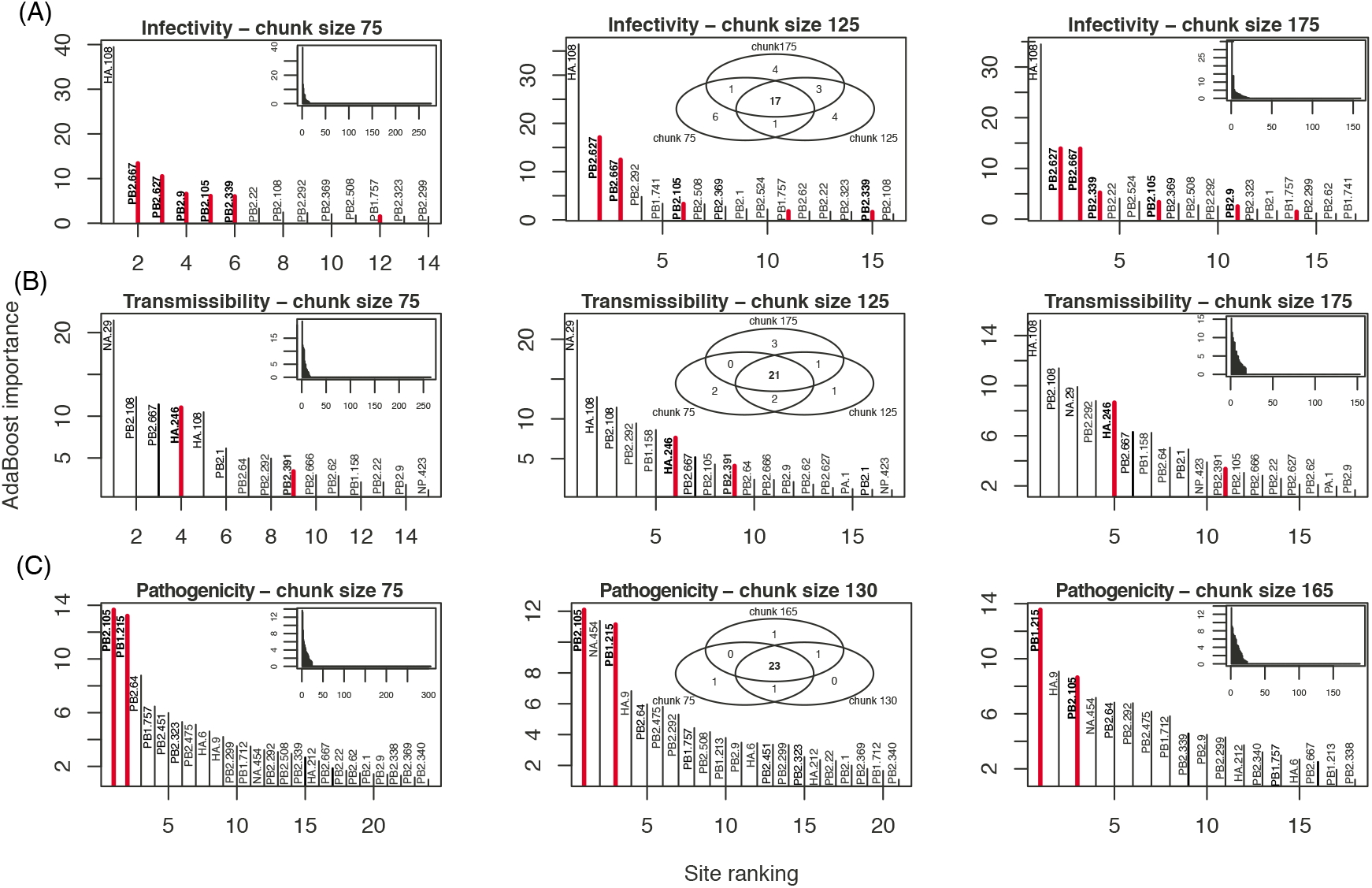
Effect of chunk size on the distribution of importance of sites for the AB algorithm. The genes and sites identified as genetic determinants of influenza phenotypes are shown for: (A) infectivity, (B) transmissibility, and (C) pathogenicity. Only results for the smallest (75 amino acids), intermediate (125), and largest (175) chunk sizes are shown. Only the most important sites (importance > 1) are shown in each panel, with sites backed by with experimental evidence highlighted in red. Insets show the whole distribution of importance values (left and right columns), and the Venn diagrams of the most important sites at all three chunk size (middle column).

One tendency that emerged when increasing chunk sizes was a decrease in sensitivity for the first pass of the algorithm, and perhaps an increase in its final specificity for both AB and RRF. In the case of infectivity for instance, the algorithms found a total of about 275 sites at chunk size 75, and roughly 150 sites at chunk size 175 (Figure 2, insets with complete distributions) for AB, and 130 sites with RRF. (Figure 3, likewise). The same pattern was observed for the two other phenotypes, transmissibility and pathogenicity. Although beyond the scope of this work, it is possible that an optimal chunk size exists in terms of area under the Receiver Operating Characteristic curve (balance between sensitivity and specificity); however, larger labeled data sets would be required for testing this possibility.

**Figure 3.**
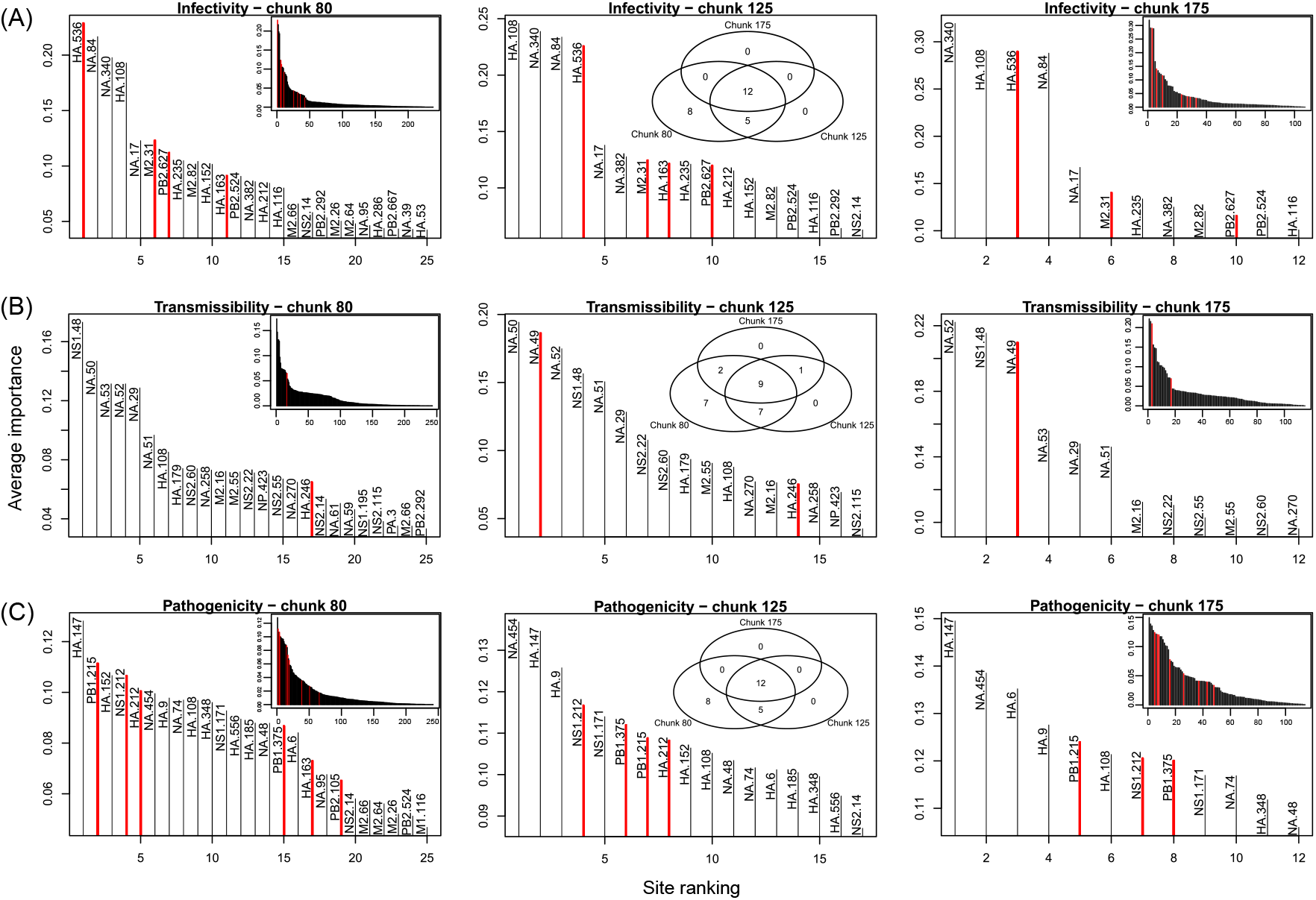
Effect of chunk size on the distribution of importance of sites for the RRF algorithm. The genes and sites identified as genetic determinants of influenza phenotypes are shown for: (A) infectivity, (B) transmissibility, and (C) pathogenicity. Only results for the smallest (80 amino acids), intermediate (125), and largest (175) chunk sizes are shown. Only the most important sites (Gini index in top 90^th^ percentile of its distribution over all the sites) are shown in each panel, with sites backed by with experimental evidence highlighted in red. Insets show the whole distribution of importance values (left and right columns), and the Venn diagrams of the most important sites at all three chunk size (middle column).

### Rediscovery of influenza determinants

So far the results showed that the predictions made by AB were less stable across chunk sizes than those from RRF, but they did not tell us anything about their accuracy. While the results in Figures 2–3 suggested that more true positive sites were found by AB (with an average across chunk sizes of 5.7, 2, and 2 for each phenotype) than by RRF (with similar averages at 3.7, 1.3, and 4.3), Table 1 suggests a more nuanced view, where RRF found 44% more sites than AB that are backed by experimental evidence (thirteen with RRF, nine with AB). Note finally that AB seemed to be better at finding sites in PB2 (the longer gene), while RRF’s rediscoveries were more evenly spread. Unfortunately, with such small numbers, the significance of these results is difficult to gauge.

Focusing on each phenotype, infectivity had the largest number of sites detected, the largest number of experimentally validated positions, and was the phenotype with the most stable predictions among the three phenotypes – see Table 1 for a full list. However, a number of sites detected by our approach, sometimes with consistent high importance values (*e.g*., HA 108 for infectivity; NA 29 for transmissibility), are, to our knowledge, not supported by any experimental evidence. As a result, it is possible that these predictions are false positives, even if absence of experimental evidence is not evidence of absence. We noted that with this small data set, cross-validation could not be performed to gauge the validity of these results: by splitting the samples ten times (equivalent to a leave-one-out resampling strategy), class imbalance increased dramatically, and the probability of drawing monomorphic alignment chunks (which lead the algorithm to fail) also increased. On the other hand, a number of key sites listed in the Influenza Research Database are also missed by our machine learning algorithms (Table 1). Furthermore, additional sites, which were not detected here, are known genetic determinants, but in other species. For instance, PB2 256 is known to increase polymerase activity, and hence boost infectivity, at least in pigs [26]. Likewise, PB2 28, 274, 526, and 607 do the same, but in birds [27]. As our alignment essentially contains sequences isolated from humans, it is possible that some of the sites we uncover are highly specific to this particular host. However, among these last five positions in PB2, only 526 was found to be polymorphic. Our results are therefore promising in that most of the known influenza sites (16 out of 20, or 80%), *i. e*. those supported by experimental evidence, were rediscovered by our algorithms.

### High sensitivity of RRF with chunking

In order to better understand the performance of the RRF algorithm, we conducted a simulation study (a similar study for AB could not be performed because of its high computational cost). For this, we generated sequence alignments on a 20-letter alphabet, representing the 20 amino acids found in the influenza proteome, with a single site whose amino acids match perfectly a binary phenotype (Figure S13). We first generated alignments under a balanced design, where half of the sequences were from the first phenotype, just as in the influenza data. The results show that for alignments with at least 30 sequences, irrespective of their sequence lengths, RRF has maximum sensitivity (Figure 4A), and high specificity (Figure 4B). With data sets containing 20 sequences or fewer, sensitivity decreases quickly as sequence length increases, but as suggested by the empirical results, smaller chunk sizes can maintain sensitivity to at least 50% at 5,000 sites. With ten sequences 5,000 sites in length, as in our influenza data set, sensitivity is pretty much 0, even for the smallest chunk size tested in our simulations (10%). Additional simulations showed that, at a chunk size of 2%, as in the influenza analysis above, sensitivity could reach 10%, but also that doubling the number of sequences in the alignment could increase sensitivity to 80% (Figure S14). As expected, class imbalance decreased both sensitivity and specificity (Figure 4C-D), which justifies why we tried to achieve a balanced distribution of phenotypes.

**Figure 4.**
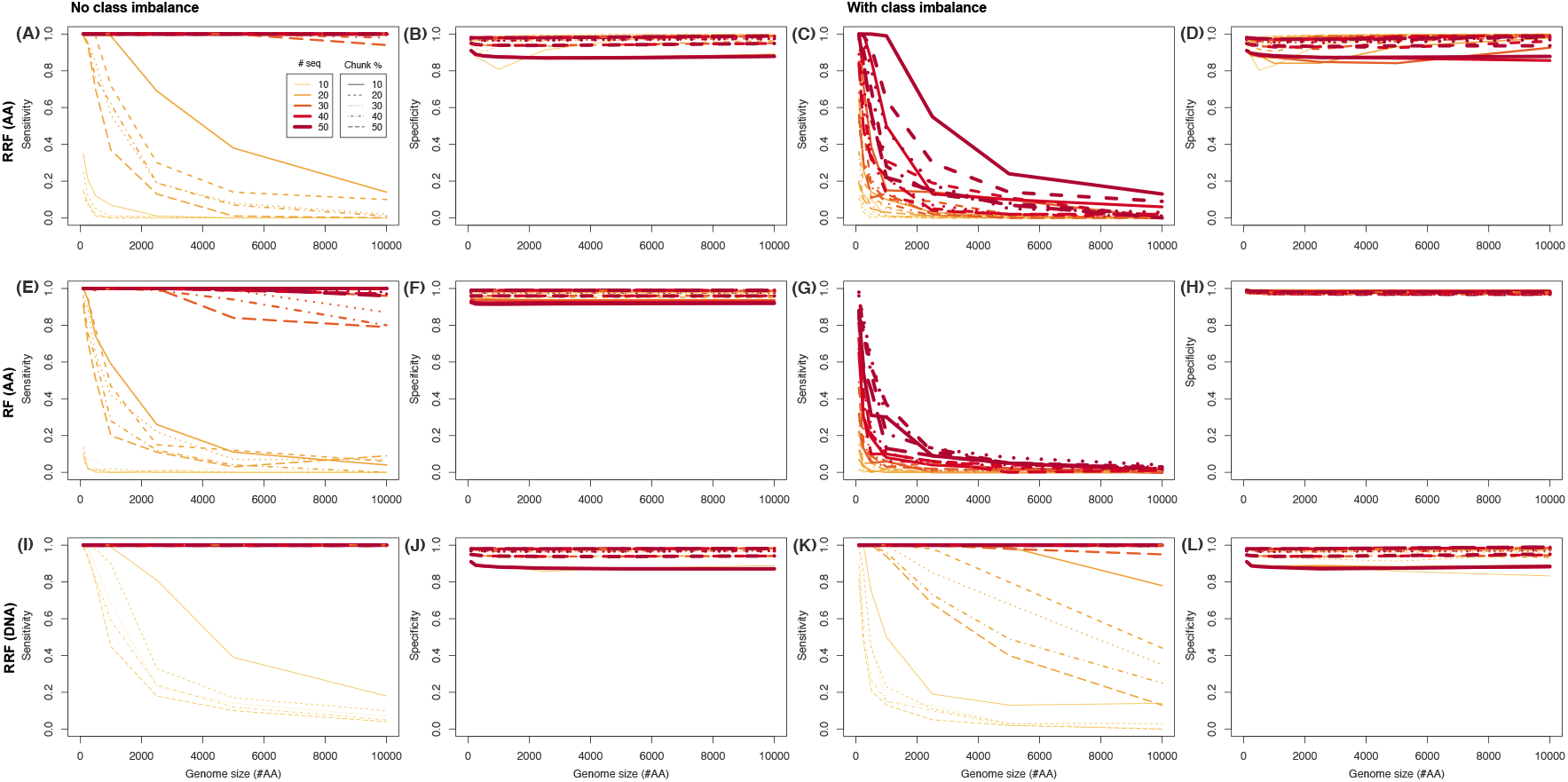
Impact of chunking and data size on sensitivity and specificity. Simulations were conducted to assess the impact of chunking, number of sequences and length of protein alignments on (A) sensitivity and (B) specificity of the RRF algorithm, with no class imbalance, or in the presence of class imbalance (C) and (D), respectively. Similar simulations were conducted under the RF algorithm (E), (F), (G) and (H), respectively, still for protein data, and under RRF for DNA data (I), (J), (K) and (L), respectively.

With such a balanced distribution, the specificity of RF was always larger than that of RRF (Figure S15). However, sensitivity of RRF was higher than that of RF (Figure 4E), and could reach 100% with as few as 20 sequences if DNA data were analyzed (Figure 4I-J), even in the presence of class imbalance (Figure 4K-L). In the balanced case, the difference between RF and RRF was highly significant (*P* ≪ 0.01), except for chunk sizes ≥ 40% (Figure S16). Again, there was a very significant interaction between chunk size and the number of sequences, with sensitivity increasing with smaller chunk sizes and larger numbers of sequences, reaching 100% with as few as 30 sequences and chunk sizes as large as 10% (Figure S16).

Altogether, our simulations are in line with the empirical results obtained on the influenza data containing few sequences, in that our true positive rates were low between 4 and 25% (Figures S14-16).

### Unimpressive performance of AB on *P. aeruginosa*

Given these encouraging simulation and empirical results on the influenza data, for which experimental evidence supported some of the identified genetic determinants of three complex phenotypes (infectivity, transmissibility, and pathogenicity) with a small number of strains (*n* = 10), we analyzed an alignment of previously sequenced bacteria (*n* = 26), for which we had access to minimum inhibitory concentration (MIC) values for three antibiotics (Ciprofloxacin, Ceftazidime, and Gentamicin) [25] – and for which simulations suggested that we could reach a sensitivity > 80% with a specificity ~ 100% (Figure 4). We first employed AB, as this algorithm performed well in a recent small sample size application [16]. For each phenotype (MIC value; Figure 5, top row), we ran the AB algorithm twice with a chunk size of 5000 (“runs A”; ~ 1% of total alignment length), and twice with a chunk size of 1000 (“runs B”; ~ 0.2% of total alignment length). This allowed us to further test how chuck size affects which sites are detected, and how stable this detection is across different chunk sizes. As with the influenza alignment, increasing chunk size tended to lower sensitivity in the first pass of the algorithm, as fewer positions of interest had high importance (Figure S17, insets). On the other hand, sensitivity seemed to be restored, in larger chunk sizes, after the second pass, as more sites with importance values > 1 were found for all phenotypes (Figure S17, main panels). Figure S18 shows that only the most important sites were identified with very similar importance values across the four runs (and hence the two chunk sizes). When only the top 25 sites were compared across these four runs, only five to seven sites were shared (Figure S19). Some of these predictions are sensible: DNA gyrase subunit B (gyrB) is known to be involved in drug resistance [28, 29], hypothetical protein PA14_40040 (see Ceftazidime phenotype runs) is known for its involvement in antibiotic biosynthesis processes [30], and tonB2 is inferred to be an iron transporter which potentially affects bacterial drug resistance [31]. Detecting genes that were previously unknown to be involved in drug resistance is not uncommon, as a recent study of antibiotic resistant *P. aeruginosa* found that 12% of the assayed strains carried novel genetic determinants [32]. What is more unexpected though is that some of these genes are not supposed to be involved in the mode of action of these specific drug: for instance, Ciprofloxacin, as a fluoroquinolone, is known to disrupt gyrB [29], but it is also the only drug in our results for which gyrB was not identified (Table 3). Likewise, sbrR is a factor involved in swimming ability [33], and would hence had been expected to be involved in the resistance to cephalosporins such as Ceftazidime, not the aminoglycoside Gentamicin (Table 3).

**Figure 5.**
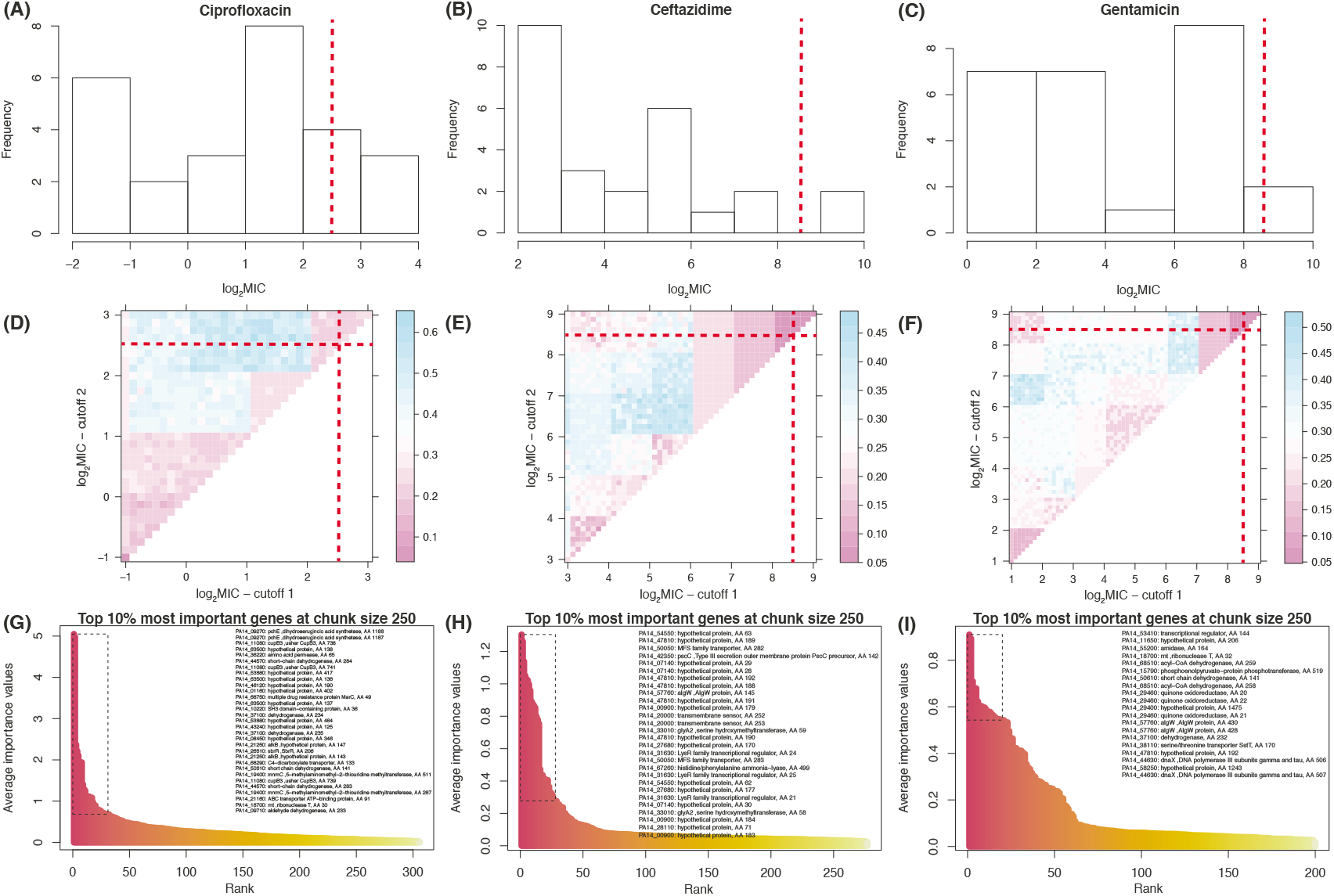
Analysis of the *P. aeruginosa* data across the 26 strains from [25]. The distributions of MIC values (on a log_2_ scale) are shown for (A) Ciprofloxacin, (B) Ceftazidime, and (C) Gentamicin. These empirical distributions were used to determine MIC thresholds for the AB analyses (Table 2). Note that the scales on the *y*-axis vary slightly. With RRF, a throughout search of the discretization was performed to select thresholds *θ*_1_ / *θ*_2_ that would minimize the Out-of-bag error for (D) Ciprofloxacin, (E) Ceftazidime, and (F) Gentamicin (color scale to the right of each panel). The *θ*_1_ / *θ*_2_ combinations (red dotted lines, also reported in first row) were determined visually. The top 10% most important sites (as per their Gini index) are highlighted (box with broken lines) among sites selected at the end of the first tier of the chunking algorithm for (G) Ciprofloxacin, (H) Ceftazidime, and (I) Gentamicin. These top sites are listed in the top right part of each distribution.

All these results were obtained with AB under one way of discretizing the MIC distributions (setting 1; Table 2). By using two other discretization schemes of the MIC distributions, it is striking that there was almost no consistency among the results (Table 3). The only proteins identified under all three settings were PA14_40040, for Ceftazidime, and tonB2 for Gentamicin. Even well-known factors such as gyrB were not identified under all three settings. Potential nonexclusive reasons for this lack of consistency include class imbalance and a rather small number of strains (*n* = 26) leading to unstable training.

**Table 2.**
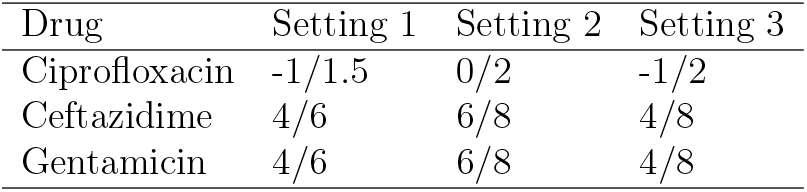
The different sets of MIC thresholds employed to assess the robustness of the classification results with AB in the case of *P. aeruginosa*. Shown are the thresholds *θ*_1_ / *θ*_2_ used on a log_2_ MIC scale: for instance, setting 1 for Ciprofloxacin means that MIC is low when ≤ −1 (*θ*_1_ = −1), high when MIC > 1.5 (*θ*_2_ = 1.5), and medium in-between.

| Drug | Setting 1 | Setting 2 | Setting 3 |
| --- | --- | --- | --- |
| Ciprofloxacin | -1/1.5 | 0/2 | -1/2 |
| Ceftazidime | 4/6 | 6/8 | 4/8 |
| Gentamicin | 4/6 | 6/8 | 4/8 |

To better quantify general performance of AB in the case of *P. aeruginosa*, we finally performed a ten-fold cross-validation (CV) analysis on the 26 strains. The confusion matrix, which depicts the predicted number of strains in each MIC category (low / medium / high) in rows, and observed numbers in columns, showed that under setting 1, class imbalance can be quite high for resistance to the three drugs, systematically leading to one of the three MIC categories with absolutely no prediction. This can be seen for instance at medium MIC values for Ceftazidime and Gentamicin (middle row):

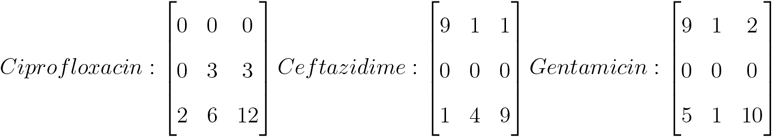

The resulting error rates for predicting the correct MIC categories were 42.31%, 30.77%, and 26.92%, respectively. As there are three discrete MIC categories, with little class imbalance, the error rate for a random classification should be close to 67%. The AB algorithm did not have a good performance, even if it still did better than chance alone at predicting the MIC category (low / high) of an unknown bacterial strain from its proteome only. Yet, these performances explained neither the instability of the pseudomonas results, nor their lack of complete biological sensibility.

One possibility is that AB was actually overfitting the data: this happens when a particular classifier accommodates all the most minute singularities of a train data set, and performs poorly on a test data set. While AB is generally considered to be robust to overfitting [34, 15], this can occur when too many iterations are performed [19] – which prompted some authors to consider *m_final_* as the AB’s main tuning parameter [35]. To assess whether overfitting was responsible for the poor pseudomonas results, we reran the AB analyses under setting 1 (Table 2) with different numbers of iterations (*m_final_* ∈ {10, 25, 50,100, 200}), both in the first and second passes of the chunking algorithm. To further assess the potential interaction with chunking, these additional analyses were run for both chunk sizes (1000 and 5000). Figure S20 shows that most analyses show a convex (concave up) CV error rate, which entails the existence of an optimal *m_final_*, that allowed optimal errors rates to be as low as 25%. Different chunk sizes had different optimal *m_final_*: for Ciprofloxacin, *e.g*., the minimum CV error rate was at *m_final_* = 50 at chunk size of 5000, but at the edge of the *m_final_* interval tested for chunk size 1000. Importantly, under both chunk sizes, the only unambiguously identified protein was trpI (PA14_00460, Table 3, underlined), a transcriptional activator implicated in Tryptophan biosynthesis. Mutations in this gene lead to reduced (albeit modest) swimming motility [36], implicated in ciprofloxacin resistance [37]. Table 3 shows that for the two other drugs, under optimal *m_final_* values, the genes identified with both chunk sizes were already among the top ten genes identified by the previous analysis, which suggests that, for a given discretization scheme of the MIC values (phenotypes), these gene lists were fairly robust to the number of iterations (*m_final_*), and that overfitting was probably not an issue in this application.

**Table 3.**
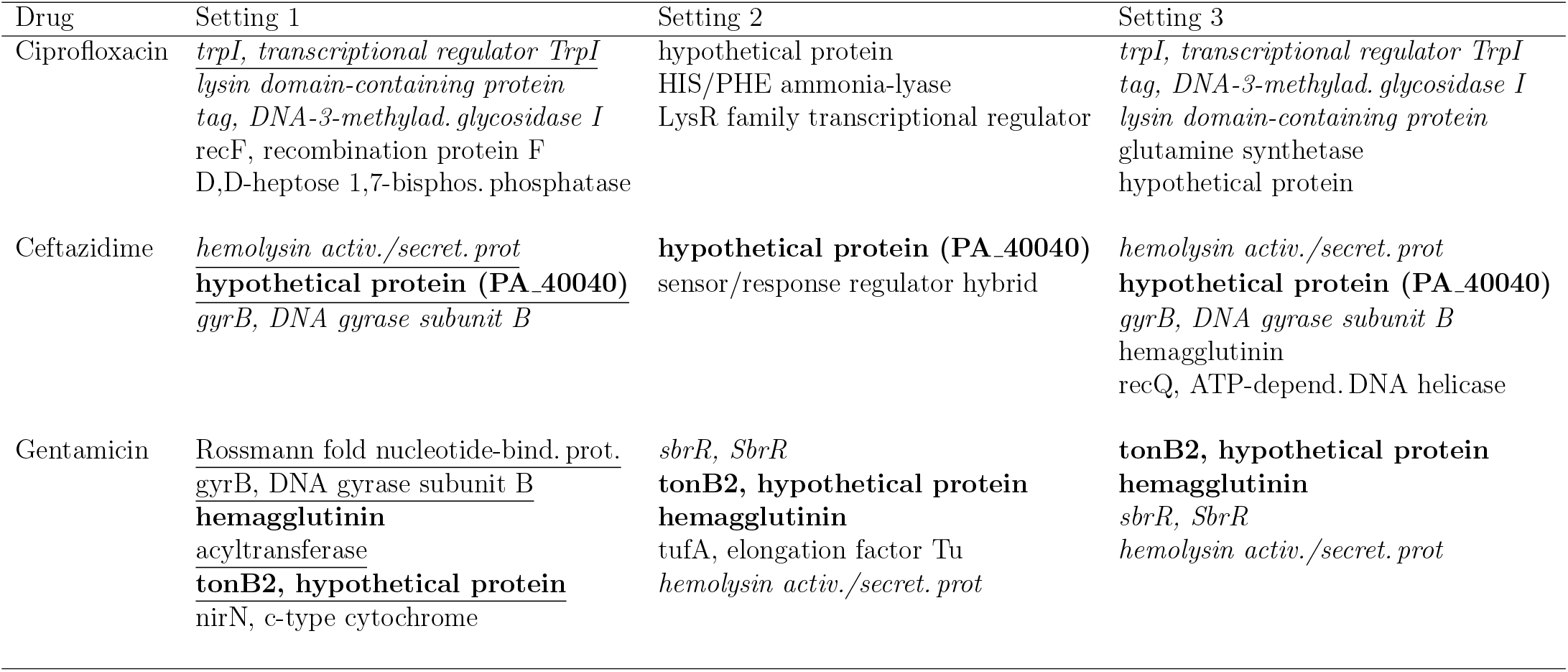
Gene lists of the most important candidates for drug resistance in *P. aeruginosa*. Shown are the genes identified in all four runs under the four settings defined in Table 2. For each drug, the genes identified in all three settings are highlighted (boldface), as well as those found in two out of the three settings (italics). Gene names that are underlined (setting 1 only) are those identified during the cross-validation experiment, under both chunk sizes.

### RRF outperforms AB on *P. aeruginosa*

AB showed some obvious shortcomings when analyzing the pseudomonas data, some of which could be linked to our discretization of the MIC curves, in addition to large memory footprints and runtimes. RRF allowed us to address all these issues: as this algorithm ran almost four orders of magnitude faster than AB (Figure 1), it was possible to perform a more thorough search of the discretization space by varying systematically the position of the two thresholds (*θ*_1_, *θ*_2_) (Figure 5A-C). The pattern of Out-of-bag error, with triangles of low errors along the diagonal (Figure 5D-F), showed that these data should be analyzed with a single MIC threshold: dissimilar (*θ*_1_,*θ*_2_) thresholds led to high errors; similar (*θ*_1_,*θ*_2_) thresholds led to low errors. This pattern was not unexpected given the coarseness of the MIC distributions (Figure 5A-C). While both Ceftazidime and Gentamicin had clear threshold minimizing the Out-of-bag error, multiple choices existed for Ciprofloxacin. To identify genetic determinants at high drug levels (the mutations conferring the highest levels of resistance), a high threshold was chosen (Figure 5A,D).

Figure 5G-I shows the most important sites identified by RRF. To determine which of these sites are potential candidates for drug resistance, we used the empirical rules derived from the analysis of influenza data, as those analyses were informed by experimental evidence, and validated by our simulations (Figure 4). As above, we expected that most of the true positive sites are (i) found in at least 50% of the repeated parts of RRF, and (ii) in the top 10% of the global distribution of importance (Gini) values. For influenza, these two rules allowed us to capture all but one of the experimentally-validated sites, while minimizing the number of false positives (see Figure S19). Under these two rules of thumb for site discovery, we found lists of sites (Figure 5) that are completely different from those found with AB (Table 3). However, these lists of sites were very stable across a wide range of chunk sizes (Figure S21), and their content made sense in light of what is known about these three drugs, which are commonly employed to treat *P. aeruginosa* infections of cystic fibrosis patients (Table 4). Note that Out-of-bag errors rates were all < 10% (Figure 5), when CV error rates under AB were at least 25%. These lists of candidate genetic determinants for drug resistance are still long, with many hypothetical proteins, and hence remain problematic for experimental validation, but large and deep mutational screens have already started to revolutionize the field [38].

**Table 4.**
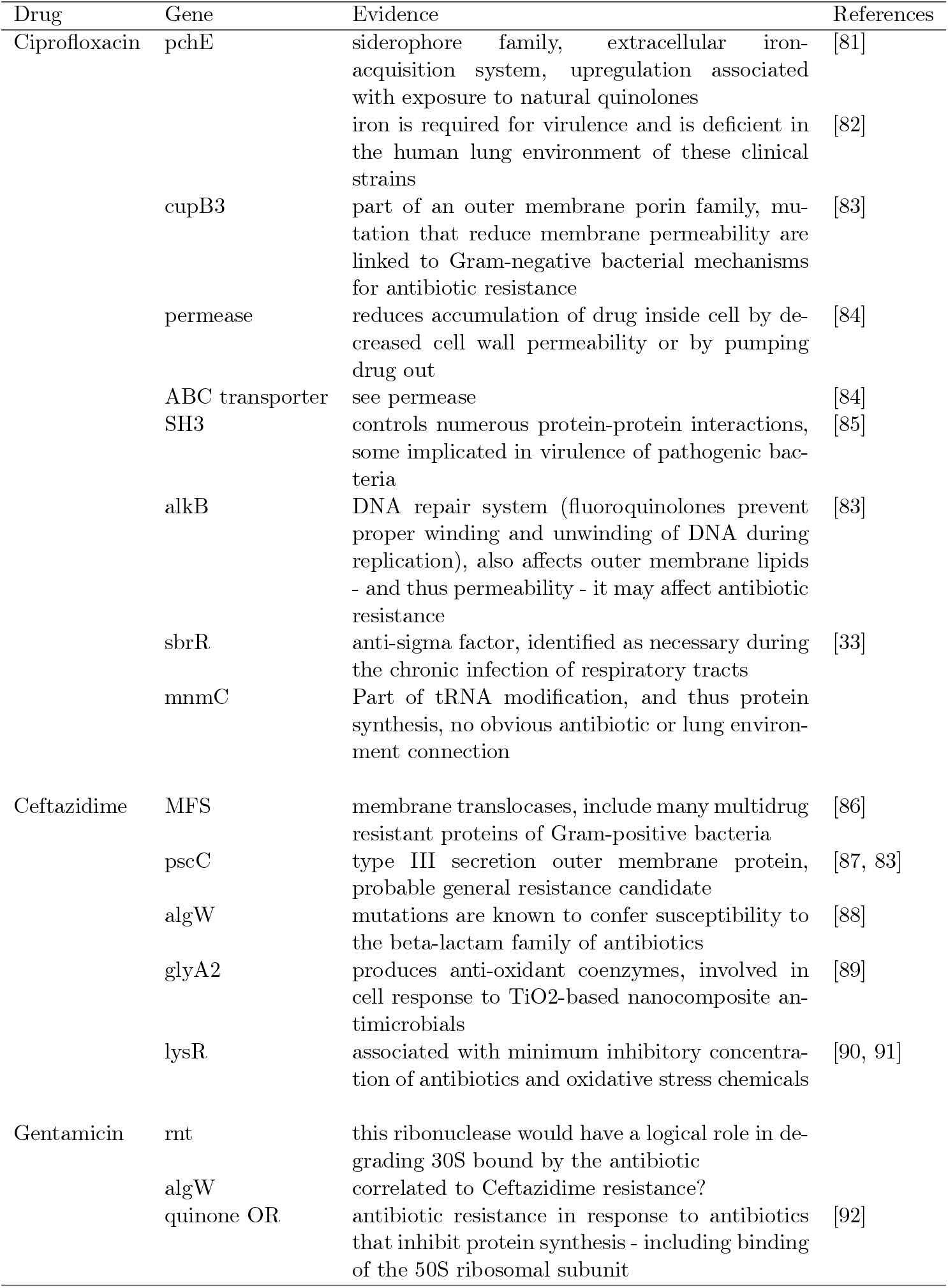
Characterization of RRF drug resistance candidates in the pseudomonas data from the literature.

Lastly, to evaluate the overall performance of RRF on the pseudomonas data set, we conducted additional simulations, as above, but with a smaller chunk size (0.2%, as in “runs B”), varying the number of sequences between 24-30, and increasing sequence lengths all the way to 10^6^ sites. Only 25 replicates were done in these conditions, but these results suggest that we would need at least 30 pseudomonas proteomes (of length 500k sites) to reach a sensitivity of 90%, and a specificity just above 60% (Figure S22). With the data set that we had access to (26 proteomes), sensitivity was < 15%. Larger proteomes, containing a million polymorphic sites or more, will require sequencing at least 30 individuals to reach sensitivity values of at least 80%.

### Conclusions

In order to find the genetic determinants of particular phenotypes whole genome or proteome data, we implemented and tested two machine learning algorithms based on adaptive boosting, which has recently proved to be quite successful when analyzing small data sets [16], and random forests. Our use of these machine learning algorithms can be characterized as *unbiased*, in that they gauge the importance of every single position in a proteome without any *a priori* assumptions – in the same way as are often characterized GWAS [4] or RNA-seq studies [39, 40, 41]. However, because each proteome contains a very large number of positions, these algorithms could not be run ‘as is’ on the entire alignment, even after removing invariant positions. In regulatory genomics, where the objective is to uncover splice junctions, such machine learning algorithms typically focus on sequence windows centered on the traits of interest, thereby reducing the feature space [42, 43]. Here, we did not consider such prior knowledge (when it existed) to pre-define features, and instead took a more agnostic approach, focusing on the entire alignment of polymorphic positions. The resulting computational burden prompted us to develop the chunking algorithm, especially in the case of AB, where the initial alignment is chopped up into smaller parts or chunks. Each algorithm, AB and RRF, is run in a first pass on each chunk, to determine a set of positions of interest, which are then collated for a second pass to rank these positions by their importance in predicting a particular phenotype. Here we showed that chunking improves runtimes, without qualitatively affecting performance, both in terms of which positions are identified, and their importance values – and may even increase sensitivity of model predictions. Although our RRF algorithm is based on random forests, it is not equivalent to this latter algorithm run on more trees because we combine the results of each repeat by taking their consensus. RRF may also be reminiscent of the iterated RF algorithm (iRF) [44], but while RRF is less sophisticated, the stability of model predictions are definitely improved.

We then showed by analyzing a data set in which the genetic determinants of complex phenotypes are known (influenza) that both AB and RRF correctly identified some positions supported by experimental evidence [45], but that the top results also included some potential false positives, and missed some known sites. Simulations suggested that excellent performance could be obtained, with sensitivity and specificity both close to 100%, but with larger data sets (> 20 sequences). The analysis of a second and larger data set (26 sequences), for which the genetic determinants of complex phenotypes are not so well known (pseudomonas) allowed us to identify positions in genes known, or predicted to be, involved in the phenotypes assayed, with more sensible results obtained with RRF than with AB. Altogether, predictions based on machine learning algorithms can allow for a quicker discovery process of the genetic determinants of complex phenotypes, but should be more thoroughly compared with traditional GWAS approaches, especially for proteomes containing more than a million polymorphic sites. Far from being restricted to finding resistance genes and mutations, our approach is amenable to predicting any kind of phenotype/genotype relationships, including disease-causing mutations. However, such a use of machine learning to determine phenotype/genotype relationships might only blossom when both phenotyping and genome sequencing costs become low enough to perform these analyses on many more individuals.

## Methods

### The influenza data

The complete proteome of *n* = 10 Influenza A strains, containing the twelve canonical genes usually found in these viruses [18], were retrieved from the Influenza Research Database [24] using their search tool based on phenotype characteristics. The retrieved strains included all viral samples with experimental evidence supporting an increase of infectivity, transmissibility, and pathogenicity, the three main phenotypes in this database (retrieved Sep 2016). The detection of a polybasic cleavage site was used as a proxy for pathogenicity; while the presence of such a site alone does not indicate an increase of pathogenicity, it is nonetheless present in highly pathogenic strains [46]. The phenotypes were encoded as binary variables, since phenotypic data were only available as a Yes / No statement. Our analyses were performed blindly, as no indication of any particular mutation was included in the strain name (Table 5). Note that our selection of strains tried to minimize class imbalance, so that both infectivity and pathogenicity have a 1:1 ratio, while transmissibility data are a bit more uneven (3:7; Table 5).

**Table 5.** List of the Influenza A strains and their associated phenotypes, as used in the training of the machine learning algorithms. For pathogenicity, polybasic cleavage was used as a proxy.

| Strain name | Infectivity | Transmissibility | Pathogenicity |
| --- | --- | --- | --- |
| A/HongKong/156/97 | No | Yes | Yes |
| A/HongKong/213/2003 | No | Yes | Yes |
| A/Indonesia/5/2005 | No | No | No |
| A/Indonesia/7/2005 | No | No | No |
| A/PuertoRico/8/34 | Yes | No | No |
| A/Swine/Indiana/1726/1988 | Yes | Yes | No |
| A/Turkey/15/2006 | No | No | No |
| A/VietNam/1203/2004 | Yes | No | Yes |
| A/VietNam/3046/2004 | Yes | No | Yes |
| A/VietNam/3062/2004 | Yes | No | Yes |

The corresponding proteomic data were downloaded from the Influenza Virus Resource [47]. We focused on amino acid data, assuming that phenotypic differences are caused by nonsynonymous mutations. Only the ten most common proteins found in all influenza strains (PB2, PB1, PA, HA, NP, NA, M1, M2, NS1, and NS2) were retrieved. This was done to ensure that the results obtained could be applied to the widest selection of strains possible. The selected viral strains were essentially from human hosts. This was done to prevent any potentially confounding factors from arising due to the different cellular targets [48], or due to the large sequence difference between avian and mammalian subtypes [49]. Strains containing only one segment (*e.g., PA/Fort Monmouth/1/47-MA(H1N1*)) were discarded.

The segments of each strain were individually aligned with MUSCLE 3.8.31 [50] to ensure accuracy, and were then concatenated into a single alignment. Any missing segment in any strain was replaced with a row of gap characters to (i) ensure proper concatenation, and (ii) prevent a mismatching of protein segments between the different influenza strains, and thus prevent distortion of the alignments. After the segment concatenation, invariant sites were removed from the alignment. This was done to reduce the computational time required for adaptive boosting, as sites without mutations do not contain any information relevant to the analysis.

### Simulations

Protein alignments were simulated based on a 20-letter alphabet by drawing amino acids with replacement from a uniform distribution. Only one site was perfectly associated with a binary phenotype. Simulations could be balanced, where each phenotype is in a 1:1 ratio, or unbalanced, where only two sequences are from the first phenotype. Number of sequences were taken in (10, 20, 30, 40, 50), the length of each alignment took values in (100, 250, 500, 1000, 2,500, 5,000, 10,000), and the chunk size changed from 10 to 50% of the total alignment length in 10% increments. One hundred replicates were performed under each condition. Sensitivity and specificity of each simulation condition were recorded, both under RRF, and RF for comparison purposes. To evaluate the impact of the size of the alphabet on performance, another round of simulations were performed on a 4-letter alphabet (RRF only), representing DNA data. These simulations allowed us to count True Positive (TP), False Negative (FN), True Negative (TN) and False Positive results, and deduce sensitivity (TP / (TP + FN)) and specificity (TN / (TN + FP)) from these.

### The pseudomonas data

The proteome alignment of *P. aeruginosa* was generated by concatenating the coding regions of the *n* =26 *P. aeruginosa* genomes previously published [25]. With a reference database of PA14 (www.pseudomonas.com; [51]), an alignment for each open reading frame (ORF) was created using an in-house pipeline that: (i) stored BLASTn 2.2.30 [52] results for each ORF of each non-PA14 genome, (ii) discarded any results with identity < 90%, (iii) assembled alignments for each ORF ensuring a genome’s sequence was used only once. This was achieved by first building a scaffold from genomic sequence with only one BLASTn result, then extending incomplete scaffolds, *i.e*., those that did not cover the full range of their respective PA14 reference sequence. Extensions were done using BLASTn results that did not correlate with higher percent identity to another incomplete scaffold nor overlapped the current scaffold by more than 30 nucleotides. Scoring of genomic ranges and overlaps was performed using Bioconductor’s function GRanges [53]). Following scaffold assembly, (iv) sequences were aligned using MUSCLE 3.8.31 [50]. Any aligned ORF with < 50% of strains having non-gap characters in at least 90% of reference sites (established via PA14 sequence) were discarded. Lastly, the remaining aligned ORFs were concatenated with the perl script `catfasta2phyml.pl` (by Johan Nylander: https://github.com/nylander/catfasta2phyml/commit/5035eb). This resulted in an alignment containing 5,944 of the 5,977 ORFs in the PA14 reference genome, and a total of 1,974,843 amino acid positions. Gene annotations were obtained from the file `UCBPP-PA14.csv` available at www.pseudomonas.com/downloads/pseudomonas/pgd_r_16_2/Pseudomonas/complete/gtf-complete.tar.gz.

Each of these 26 strains had previously been characterized phenotypically, with respect to their antibiotic resistance to three different drugs: Ciprofloxacin, Ceftazidime, and Gentamicin [25]. All three are broad-spectrum drugs, used to treat patients infected by *P. aeruginosa*, and all three belong to different families of antibiotics (Ciprofloxacin is a fluoroquinolone, Ceftazidime is a cephalosporin, and Gentamicin is an aminoglycoside). As such, each drug has a different mode of action: fluoroquinolones inhibit enzymes such as DNA gyrase and topoisomerase IV, involved in the replication of DNA [29]; cephalosporins interrupt the synthesis of the peptidoglycan layer forming the bacterial cell wall [54]; aminoglycosides bind to the 30s ribosomal subunit and inhibit protein synthesis [55]. Hence, different genes can be expected to be involved in the resistance to these antibiotics. Resistance had been quantified by means of MIC assays, where a growth medium containing antibiotics is serially diluted, in two-fold steps, before an equal volume of overnight bacterial culture be inoculated into each dilution. After at least 16 hours of growth in these conditions, the MIC is defined as the minimum antibiotic concentration that does not permit growth.

### Predictive modeling

Two machine learning algorithms were used to construct a model to predict each phenotype from the proteomic information (proteome) in both data sets, influenza and pseudomonas. The first was AB [12], as implemented in the R package adabag [56], while the second was the RF [17] algorithm from the R package randomForest 4.6-14 [57]. All scripts were run in R 3.5 [58], and are available from https://github.com/sarisbro, alongside the data used.

The features included in both machine learning algorithms were the same: the amino acids of each proteome alignment. To keep track of site identity, each alignment was stored as a matrix, where column names contained the name of each ORF and the amino acid position within each ORF. As only polymorphic positions in the alignments are potentially informative, invariant sites were first discarded. This left 4,392 polymorphic positions in the influenza alignment, and 511,780 in the *P. aeruginosa* alignment.

While the AB algorithm used was unaltered, with the total number of iterations left to its default value (*m_final_* = number of sites in the alignment), we slightly modified the RF algorithm. Indeed, due to the stochastic nature of this algorithm, RF can lead to different rankings of the most important features across different runs of the same model on the same data. To alleviate this issue, we ran each random forest model ten times, keep only sites that have a Gini index > 0.075. These ten sets of features are then combined by (i) taking their consensus at a certain threshold, and (ii) keep only the top 10% consensus features (Figure S1). A similar modification of AB could have been attempted, but was not pursued here due to large memory footprint and runtimes of this algorithm.

Indeed, one limitation of most machine learning algorithms is that they can require a large amount of memory to run, especially in the case of data sets with large numbers of features, such as with the *P. aeruginosa* alignment. To alleviate this issue, alignments were split into sequential chunks of pre-specified sizes, ranging from 75 to 175 amino acids (by increments of five) for the influenza data, and chunk sizes being either 1000 or 5000 (AB), or ranging from 80 to 4000 (RRF) amino acids for the pseudomonas data. Note that these splits are largely random as segments of the influenza genome were “randomly” concatenated (by convention, segments are ordered by decreasing length, just like chromosomes in Eukaryotes), and protein-coding genes in pseudomonas were “randomly” concatenated (based on their order in the PA14 strain). All the computing in this step can be parallelized, so that each classifier can be run on each chunk independently by distributing the analyses over eight threads with the R foreach package 1.4.4 [59]. The RRF algorithm was first run on each chunk during the first pass of the chunking algorithm, hereby producing a set of positions of interest. Only those with a Gini index > 0.075 were kept (first pass of the chunking algorithm), collated, and the classifier was run a second time on these (i.e., during pass 2 of the chunking algorithm), to produce the final set of most important sites (*i.e*., predictors of each phenotype in the top 10% Gini indices in the second pass of the chunking algorithm).

The performance of the machine learning algorithms was assessed using ten-fold crossvalidations (AB) and Out-of-bag errors (RRF). For cross-validations, the alignment was divided into ten sets of sequences (samples), nine sets being used for training and the remaining one for testing. That process was then repeated for all ten subsets [56]. In the context of our chunking procedure, cross-validation was performed on the second pass of the AB algorithm. Out-of-bag errors were computed on the predictions based on the bootstrapped trees that were not included during training.

As the evolution of antimicrobial resistance in *P. aeruginosa* can include many understudied sites [60], only the influenza data have sufficient experimental validations to which we can compare our predictions and thus determine their accuracy. In order to learn from this data set how to best balance true positives (sites that are known to be experimentally validated and are detected) and false positives (sites that are detected but are not experimentally validated) from the distribution of their importance values, we defined two thresholds (Figure S1). First, the expectation is that the most important sites will mostly have true positives, so the first threshold used the influenza data to determine what top percentile of ranked importance values maximizes the number of true positive. We refer to this threshold as the “percentile threshold.” Then, to minimize the number of false positives among these top sites based on the repeated nature of the RRF, we implemented a second threshold, the “consensus threshold,” which works as follows: all replicates of the RRF were run independently, and only sites that were predicted at a certain percentile threshold (*i.e*., a consensus) of all the runs were logged. Among these, only those that were found, say in 90% of the runs (9 runs out of 10), were considered as “genetic determinants.” Intuitively, the higher this consensus threshold, the lower the number of false positives. We then varied both threshold on the influenza data to (i) maximize the number of true positives (percentile threshold) while (ii) minimizing the number of false positive (consensus threshold). We employed these thresholds to identify genetic determinants in the pseudomonas data.

Finally for the pseudomonas data, one additional step was required. As the algorithms used to identify the genetic determinants of phenotypes require categorical data, MIC distributions were discretized. To do so, and in the case of AB first, the distribution of log_2_ MIC of each drug was first plotted, and appeared to be trimodal; hence, it seemed natural to design a classification with three categories: ‘low’, ‘medium’, and ‘high’ MIC. The boundaries between these categories were determined to minimize class imbalance, and hence guarantee that each discrete category had similar numbers of samples. Because of the relative subjectivity in determining these categories, three different sets of MIC thresholds were employed to assess the robustness of the results. All analyses were run four times to further assess robustness, and stability of which sites were identified. However, when doing so, the number of classification categories goes from two (influenza) to three (pseudomonas), without any statistical justification. To address this point, a more thorough search was performed using RRF – as RRF is much faster than AB. For this, the range of MIC values for each drug was discretized into bins of width 0.125 (on a log_2_ scale of MIC values), and two thresholds (*θ*_1_, *θ*_2_) were defined, hereby dividing the distribution of MIC values into three domains. An initial classifier was then run for all combination of thresholds with *θ*_1_ < *θ*_2_, and classification errors (RRF: Out-of-bag errors) were logged, and used to define MIC thresholds to perform the final analyses. A small chunk size (100) and a regular RF algorithm was used to speed up these computations (see Results).

## Declarations

### Ethics approval and consent to participate

Not applicable. We have no human or animal data involved.

### Consent for publication

Not applicable.

### Availability of data and materials

The data and source code used in this study are available from github.com/sarisbro.

### Competing interests

The authors declare that they have no competing interests.

### Funding

This work was supported by the University of Ottawa’s Undergraduate Research Opportunity Program (GSL, MH) and assistantships (JD), and the Natural Sciences Research Council of Canada (SAB). Funding sources played no role in the design, results, or interpretations presented in this work.

### Authors’ contributions

SAB conceived the research; GSL, MH, JD and SAB wrote parts of the R programming code; JD participated in method design and data handling; GSL and MH performed the data analyses. All authors wrote parts and edited the complete manuscript, before reading and approving the final manuscript.

## Acknowledgements

We thank Jeremy Dettman and Rees Kassen for sharing with us both their MIC and sequencing data, as well as the Center for Advanced Computing and Compute Ontario for providing us access to their servers. We are also grateful to Berthin Biyong, Graham Colby, Jeremy Dettman, Rees Kassen, and Matti Ruuskanen for discussions, and to two anonymous reviewers who helped improve this work.

## Abbreviations

AB: adaptive boosting
RF: random forest
RRF: repeated random forest
ML: machine learning
GWAS: genome-wide association study
SNP: single nucleotide polymorphism
ANCOVA: analysis of covariance
MIC: minimum inhibitory concentration
CV: cross-validation

