## Supplementary Information for "Identifying genetic determinants of complex phenotypes from whole genome sequence data"

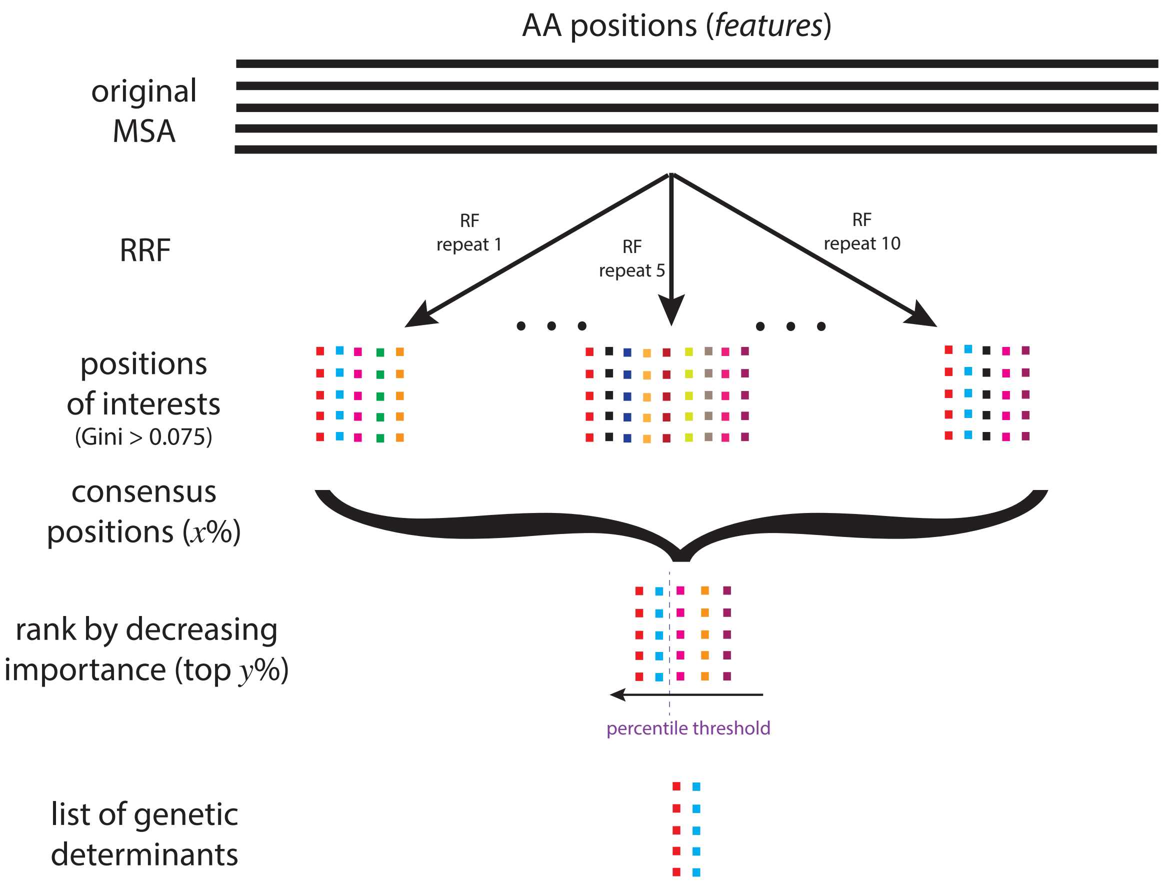


**Fig.S1.** The consensus and the percentile thresholds for the RRF algorithm. The standard random forest (RF) algorithm is run ten times on the multiple sequence alignment (MSA; in practice, each chunk), from which ten sets of "positions of interest" are deduced (with a Gini index > 0.075). Only the sites that are found in at least $x$% (for illustration purposes, $x = 50\%$ here) of the ten repeated RF runs are kept for inference according to the *consensus threshold*. These consensus positions of interest are ranked by decreasing importance value, based on the Gini index. Only the top $y$% are kept (for illustration purposes, $x = 40\%$ here) according to the *percentile threshold*.

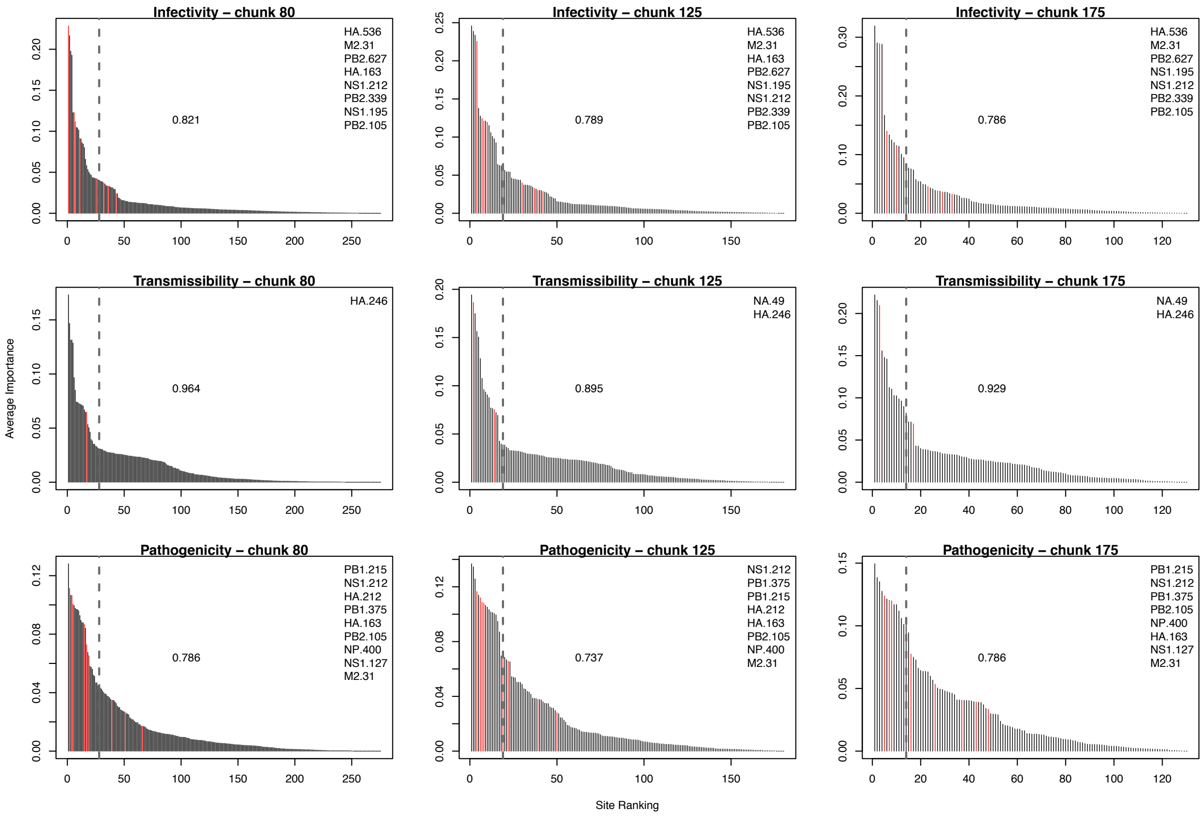


**Fig.S2.** Effect of chunk size, consensus and percentile thresholds on RRF predictions. The genes and sites identified as genetic determinants of influenza phenotypes are at a 10% consensus threshold shown for: (A) infectivity, (B) transmissibility, and (C) pathogenicity. Only the smallest (80 amino acids), intermediate (125), and largest (175) chunk sizes are shown. All the most important (Gini) sites are shown in each panel, with sites backed by with experimental evidence highlighted in red. Insets show the lists of these sites backed by experimental evidence. In each panel, the top 10% important sites are to the left of the dotted vertical line. The center of each panel shows the proportion of false positives in the top 10% important sites.

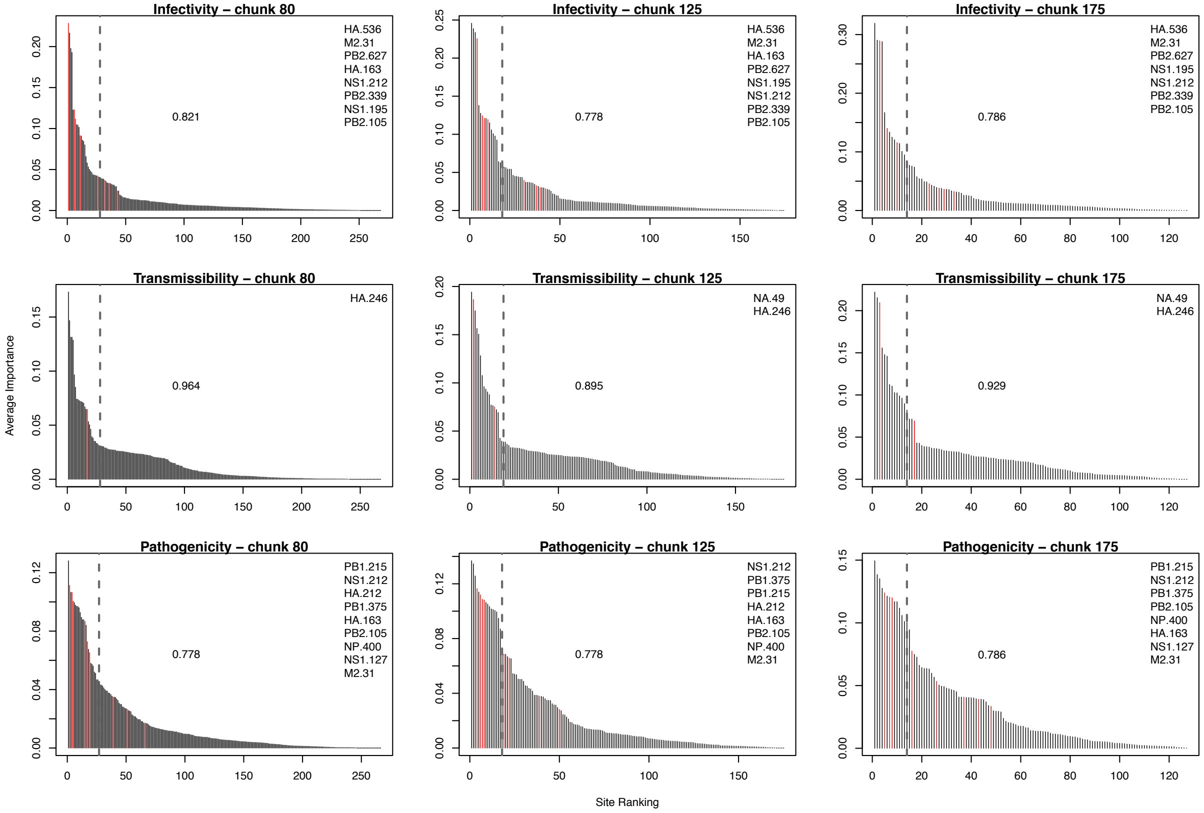


**Fig.S3.** Effect of chunk size, consensus and percentile thresholds on RRF predictions. The genes and sites identified as genetic determinants of influenza phenotypes are at a 20% consensus threshold shown for: (A) infectivity, (B) transmissibility, and (C) pathogenicity. Only the smallest (80 amino acids), intermediate (125), and largest (175) chunk sizes are shown. All the most important (Gini) sites are shown in each panel, with sites backed by with experimental evidence highlighted in red. Insets show the lists of these sites backed by experimental evidence. In each panel, the top 10% important sites are to the left of the dotted vertical line. The center of each panel shows the proportion of false positives in the top 10% important sites.

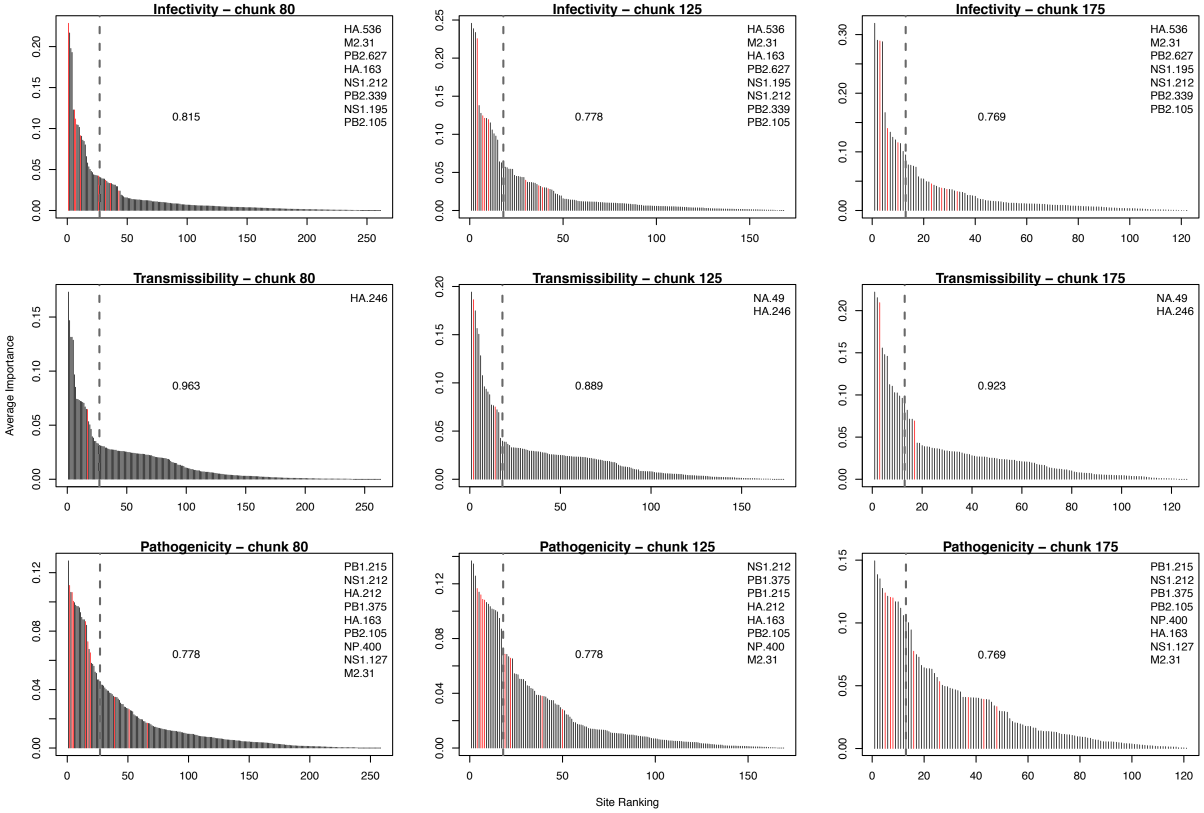


**Fig.S4.** Effect of chunk size, consensus and percentile thresholds on RRF predictions. The genes and sites identified as genetic determinants of influenza phenotypes are at a 30% consensus threshold shown for: (A) infectivity, (B) transmissibility, and (C) pathogenicity. Only the smallest (80 amino acids), intermediate (125), and largest (175) chunk sizes are shown. All the most important (Gini) sites are shown in each panel, with sites backed by with experimental evidence highlighted in red. Insets show the lists of these sites backed by experimental evidence. In each panel, the top 10% important sites are to the left of the dotted vertical line. The center of each panel shows the proportion of false positives in the top 10% important sites.

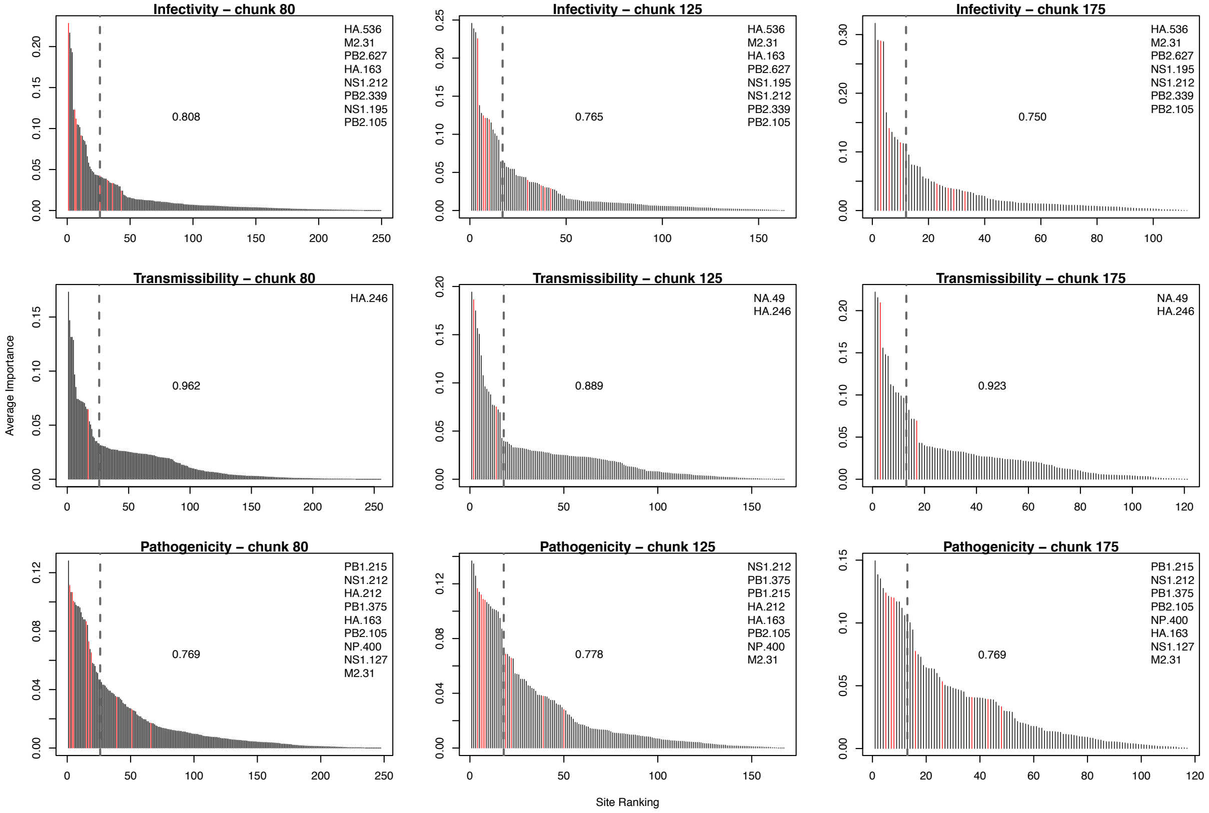


**Fig.S5.** Effect of chunk size, consensus and percentile thresholds on RRF predictions. The genes and sites identified as genetic determinants of influenza phenotypes are at a 40% consensus threshold shown for: (A) infectivity, (B) transmissibility, and (C) pathogenicity. Only the smallest (80 amino acids), intermediate (125), and largest (175) chunk sizes are shown. All the most important (Gini) sites are shown in each panel, with sites backed by with experimental evidence highlighted in red. Insets show the lists of these sites backed by experimental evidence. In each panel, the top 10% important sites are to the left of the dotted vertical line. The center of each panel shows the proportion of false positives in the top 10% important sites.

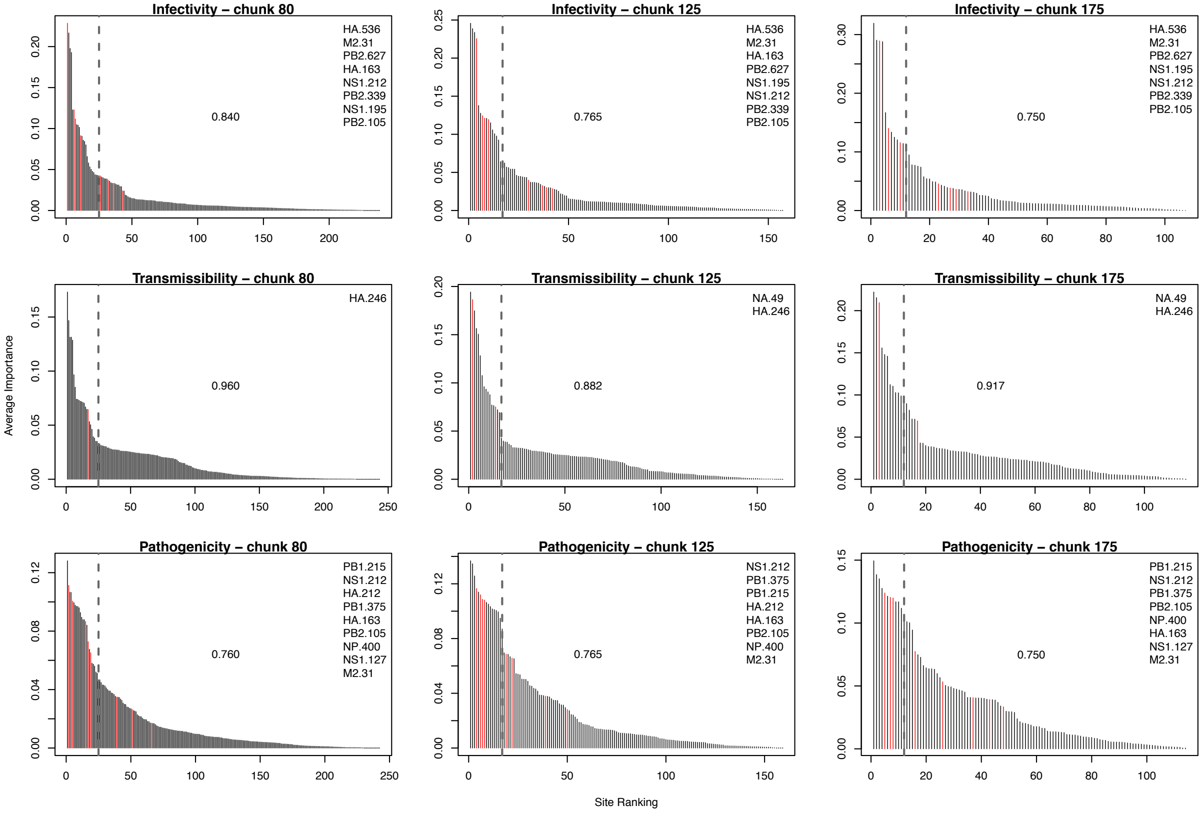


**Fig.S6.** Effect of chunk size, consensus and percentile thresholds on RRF predictions. The genes and sites identified as genetic determinants of influenza phenotypes are at a 50% consensus threshold shown for: (A) infectivity, (B) transmissibility, and (C) pathogenicity. Only the smallest (80 amino acids), intermediate (125), and largest (175) chunk sizes are shown. All the most important (Gini) sites are shown in each panel, with sites backed by with experimental evidence highlighted in red. Insets show the lists of these sites backed by experimental evidence. In each panel, the top 10% important sites are to the left of the dotted vertical line. The center of each panel shows the proportion of false positives in the top 10% important sites.

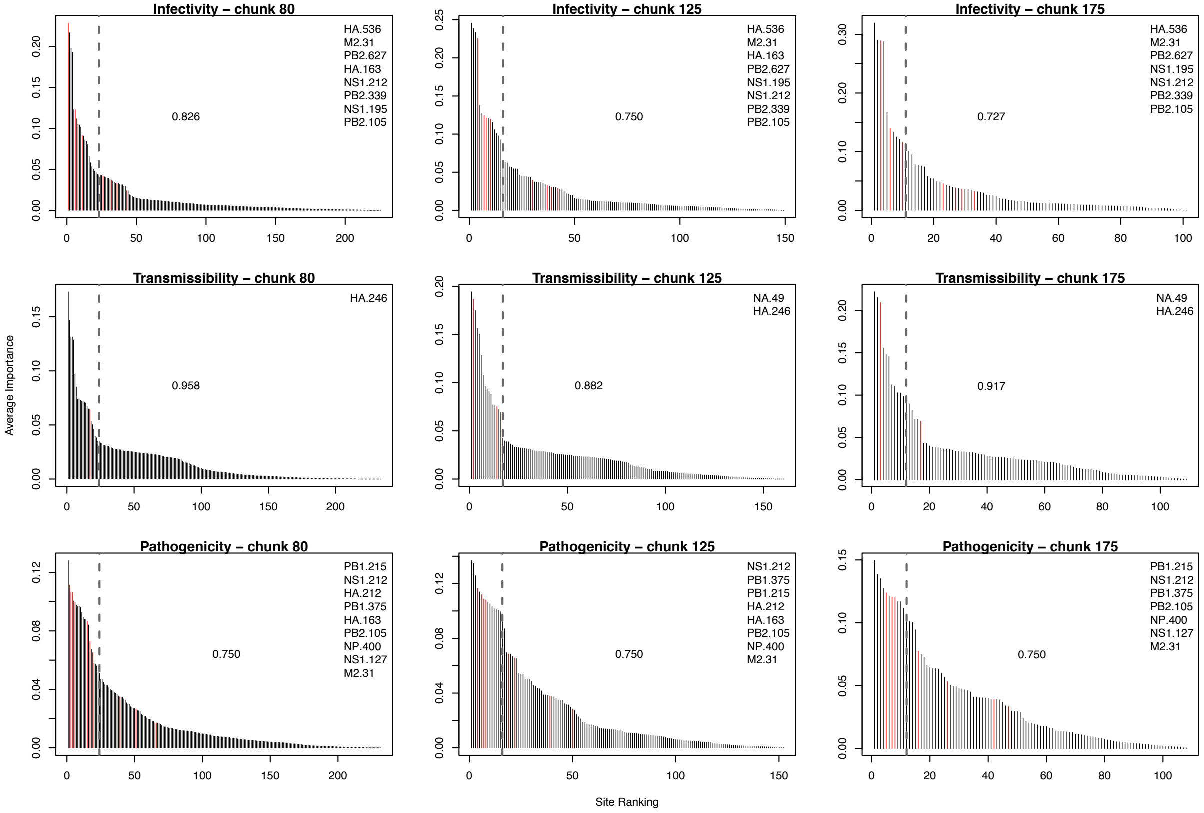


**Fig.S7.** Effect of chunk size, consensus and percentile thresholds on RRF predictions. The genes and sites identified as genetic determinants of influenza phenotypes are at a 60% consensus threshold shown for: (A) infectivity, (B) transmissibility, and (C) pathogenicity. Only the smallest (80 amino acids), intermediate (125), and largest (175) chunk sizes are shown. All the most important (Gini) sites are shown in each panel, with sites backed by with experimental evidence highlighted in red. Insets show the lists of these sites backed by experimental evidence. In each panel, the top 10% important sites are to the left of the dotted vertical line. The center of each panel shows the proportion of false positives in the top 10% important sites.

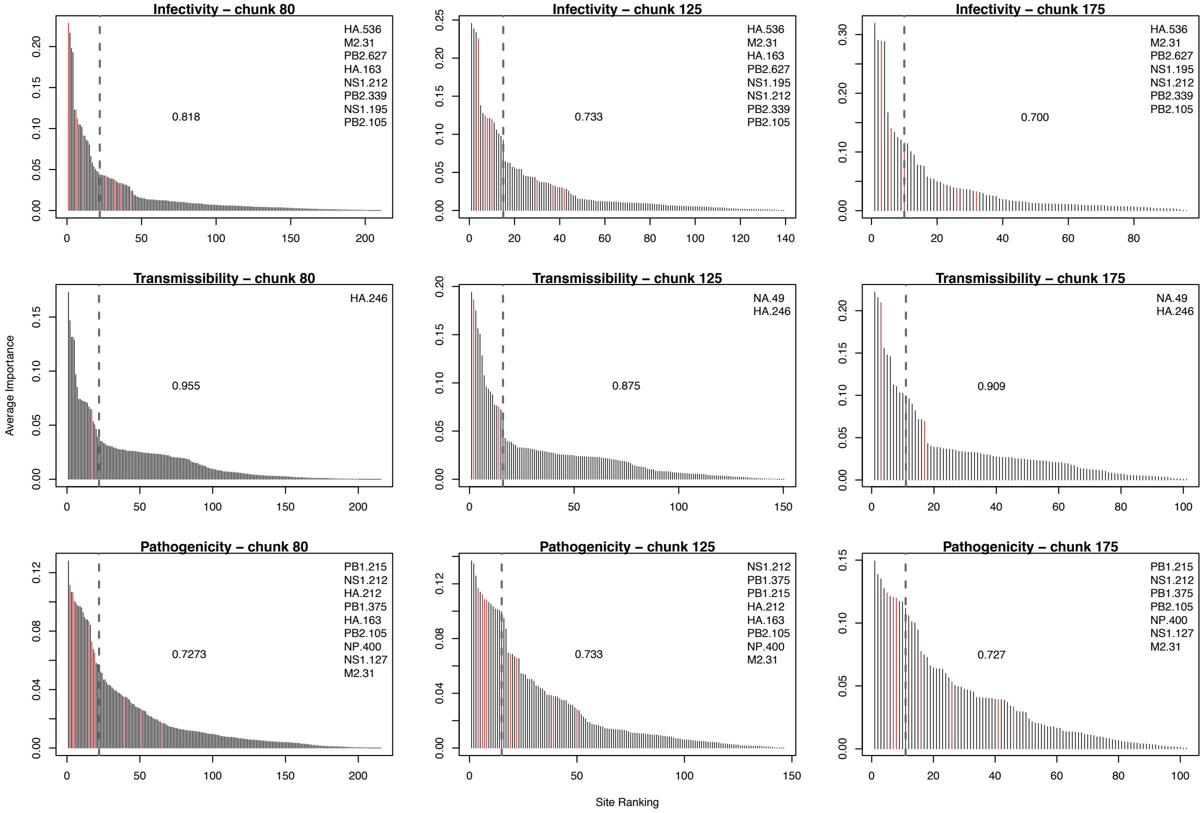


**Fig.S8.** Effect of chunk size, consensus and percentile thresholds on RRF predictions. The genes and sites identified as genetic determinants of influenza phenotypes are at a 70% consensus threshold shown for: (A) infectivity, (B) transmissibility, and (C) pathogenicity. Only the smallest (80 amino acids), intermediate (125), and largest (175) chunk sizes are shown. All the most important (Gini) sites are shown in each panel, with sites backed by with experimental evidence highlighted in red. Insets show the lists of these sites backed by experimental evidence. In each panel, the top 10% important sites are to the left of the dotted vertical line. The center of each panel shows the proportion of false positives in the top 10% important sites.

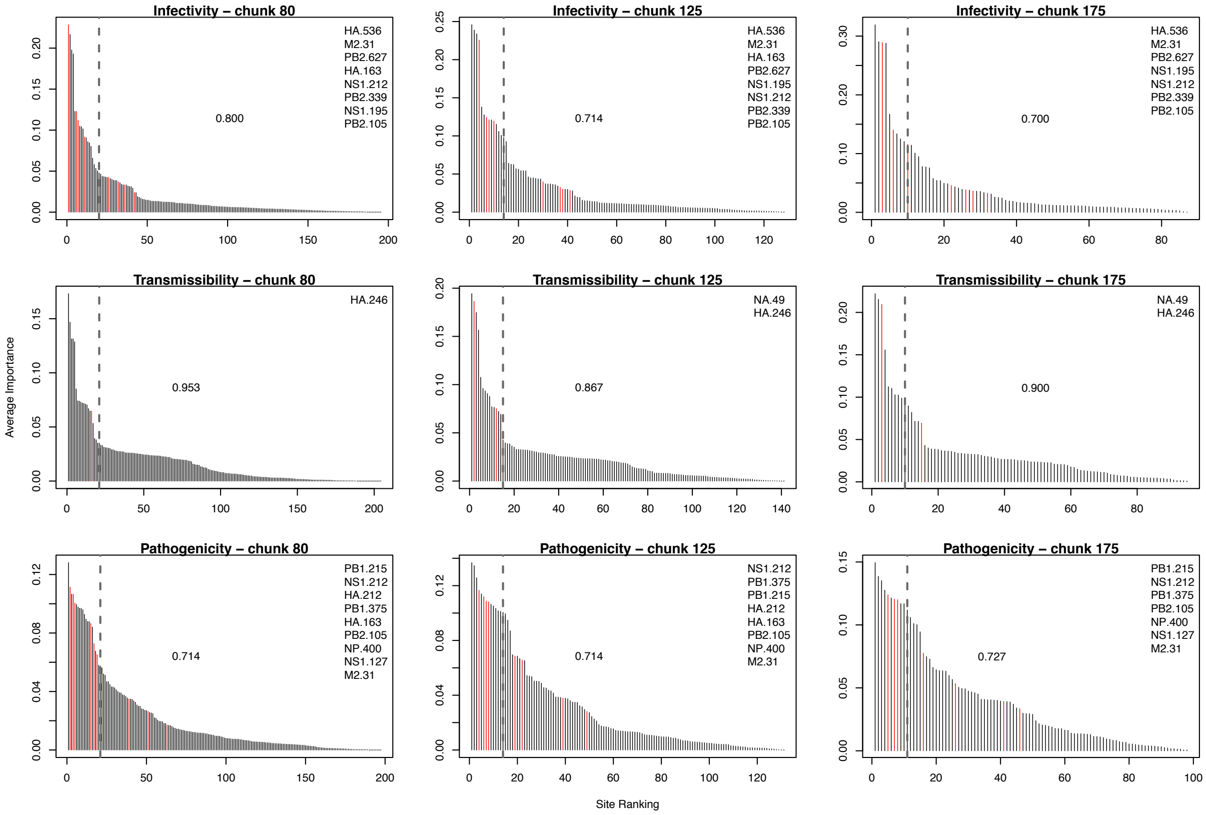


**Fig.S9.** Effect of chunk size, consensus and percentile thresholds on RRF predictions. The genes and sites identified as genetic determinants of influenza phenotypes are at a 80% consensus threshold shown for: (A) infectivity, (B) transmissibility, and (C) pathogenicity. Only the smallest (80 amino acids), intermediate (125), and largest (175) chunk sizes are shown. All the most important (Gini) sites are shown in each panel, with sites backed by with experimental evidence highlighted in red. Insets show the lists of these sites backed by experimental evidence. In each panel, the top 10% important sites are to the left of the dotted vertical line. The center of each panel shows the proportion of false positives in the top 10% important sites.

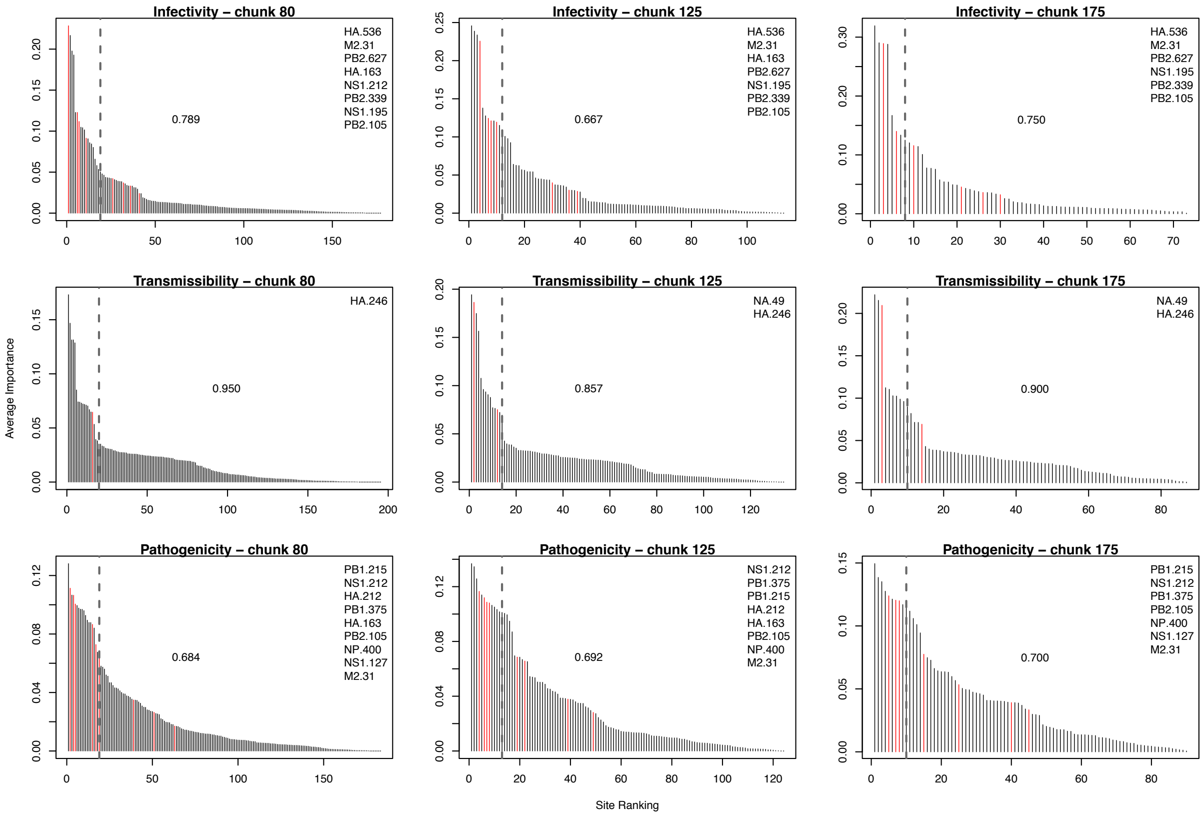


**Fig.S10.** Effect of chunk size, consensus and percentile thresholds on RRF predictions. The genes and sites identified as genetic determinants of influenza phenotypes are at a 90% consensus threshold shown for: (A) infectivity, (B) transmissibility, and (C) pathogenicity. Only the smallest (80 amino acids), intermediate (125), and largest (175) chunk sizes are shown. All the most important (Gini) sites are shown in each panel, with sites backed by with experimental evidence highlighted in red. Insets show the lists of these sites backed by experimental evidence. In each panel, the top 10% important sites are to the left of the dotted vertical line. The center of each panel shows the proportion of false positives in the top 10% important sites.

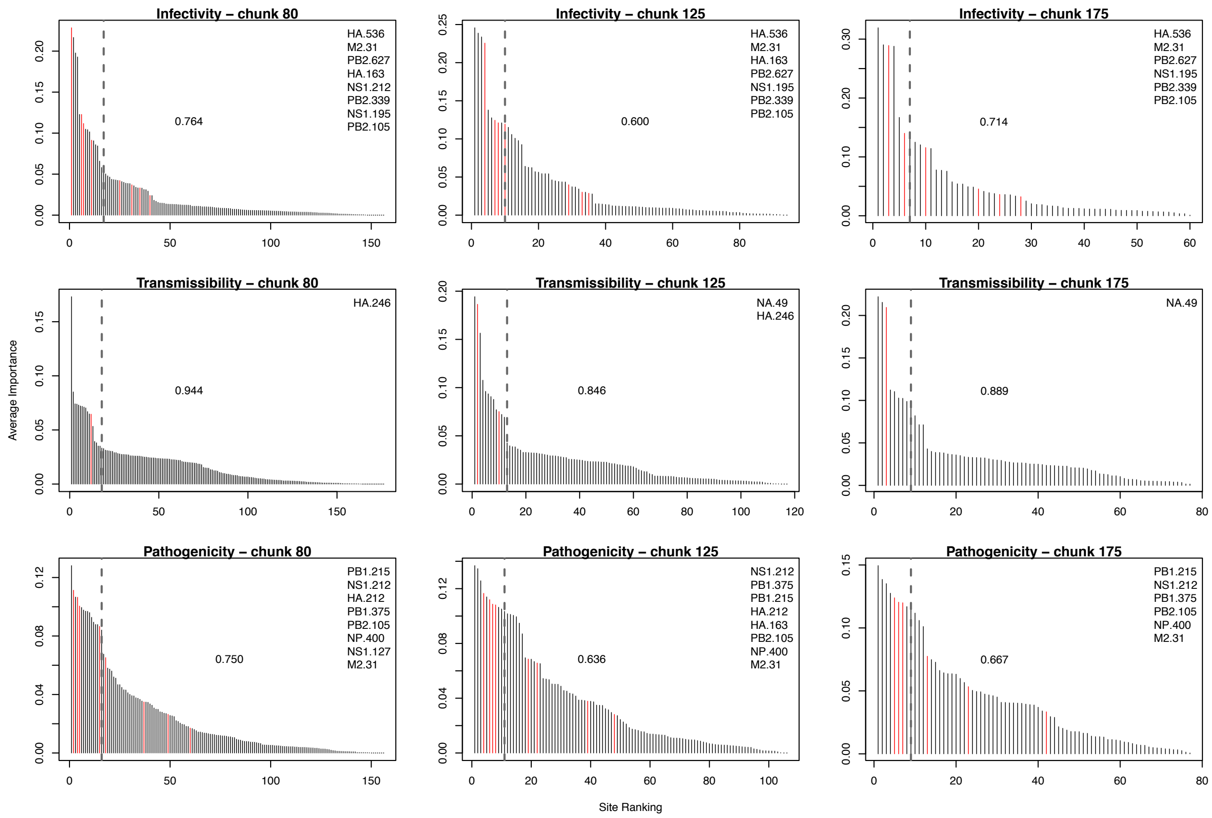


**Fig.S11.** Effect of chunk size, consensus and percentile thresholds on RRF predictions. The genes and sites identified as genetic determinants of influenza phenotypes are at a 100% consensus threshold shown for: (A) infectivity, (B) transmissibility, and (C) pathogenicity. Only the smallest (80 amino acids), intermediate (125), and largest (175) chunk sizes are shown. All the most important (Gini) sites are shown in each panel, with sites backed by with experimental evidence highlighted in red. Insets show the lists of these sites backed by experimental evidence. In each panel, the top 10% important sites are to the left of the dotted vertical line. The center of each panel shows the proportion of false positives in the top 10% important sites.

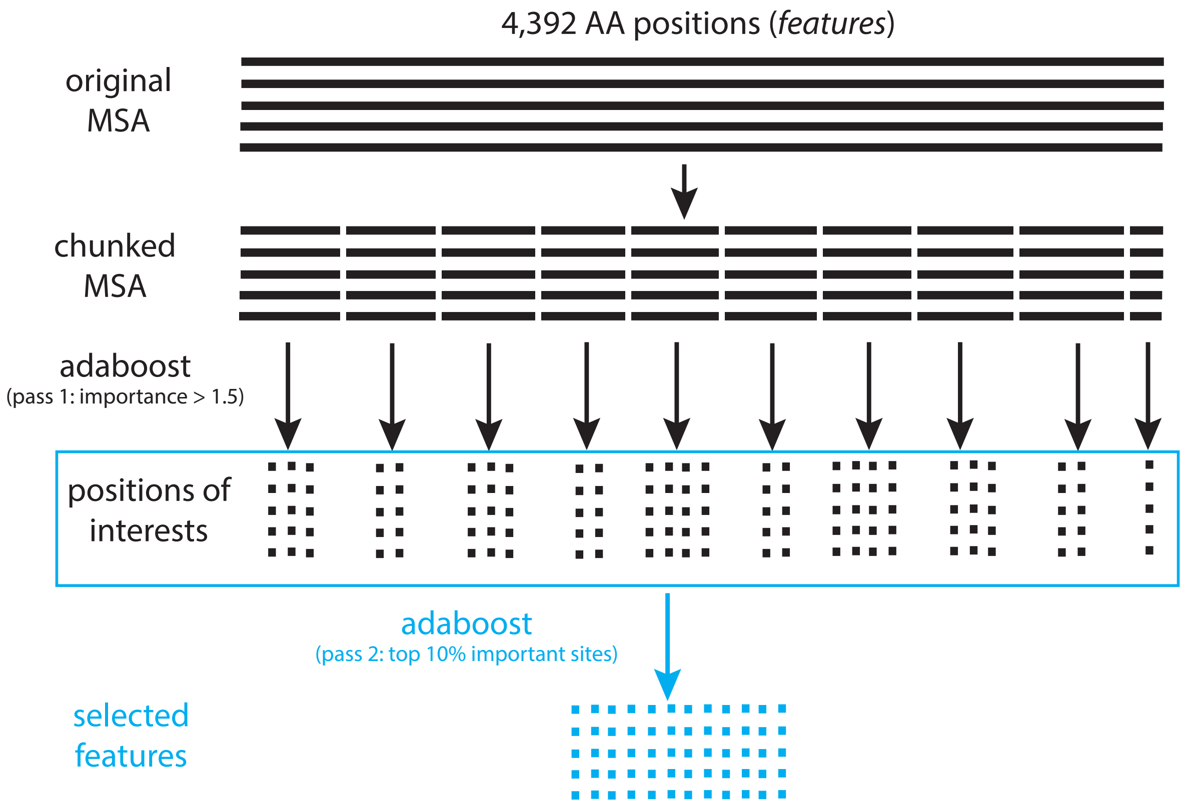


**Fig.S12.** The chunking algorithm. Illustrated in the case of the AB algorithm, each multiple sequence alignment (MSA) is subdivided into smaller parts (*chunks*), on which the ML algorithm is run in a first pass ('pass 1') to determine a set of positions of interests. A second pass ('pass 2') is run with the same ML algorithm on the entire set of positions of interests to determine the selected features over the entire MSA. In the case of RRF, the ten repeats are run on each chunk, during pass 1, while the standard RF algorithm is run during pass 2.

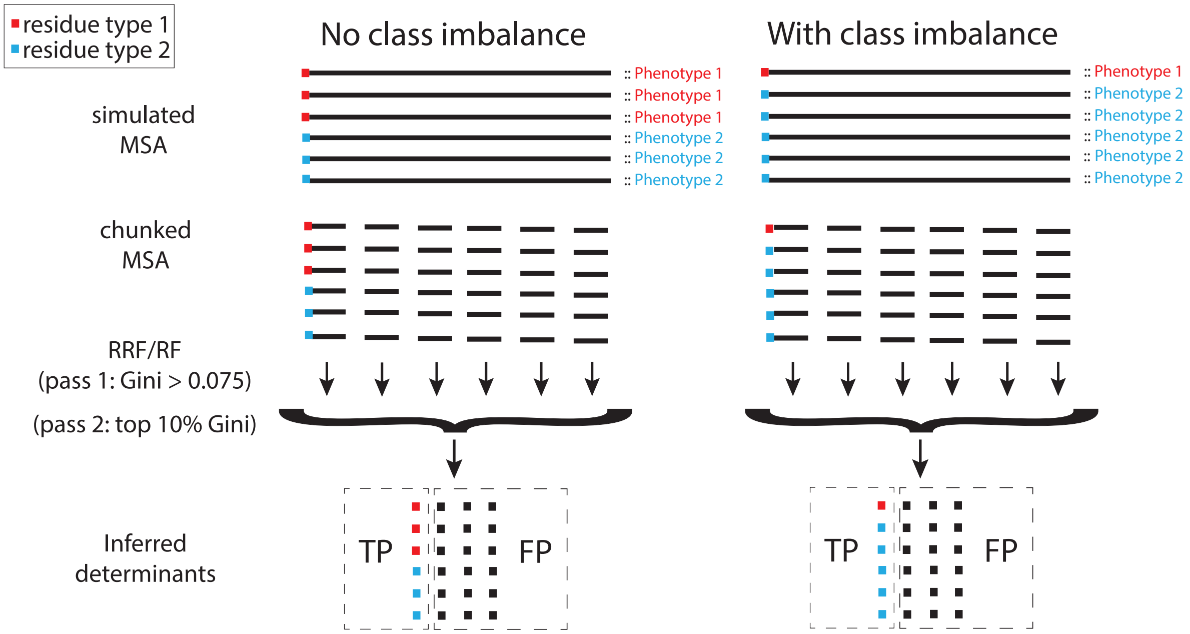


**Fig.S13.** Design of the simulation experiment. Sequence alignments were generated under two conditions, without or with class imbalance. In each case, a single position in the alignment had residues that matched perfectly the binary phenotype (in color, red and blue), while the other positions (in black) had residues drawn from a uniform distribution on the alphabet of interest (animo acids: 20 letters; DNA: 4 letters). Simulated data were analyzed under both the RRF and the RF algorithms, varying chunk size, so that True Positives (TP; shown), True Negatives (TN), False Positives (FP; shown) and False Negatives (FN) could be counted, and used to calculate sensitivity and specificity.

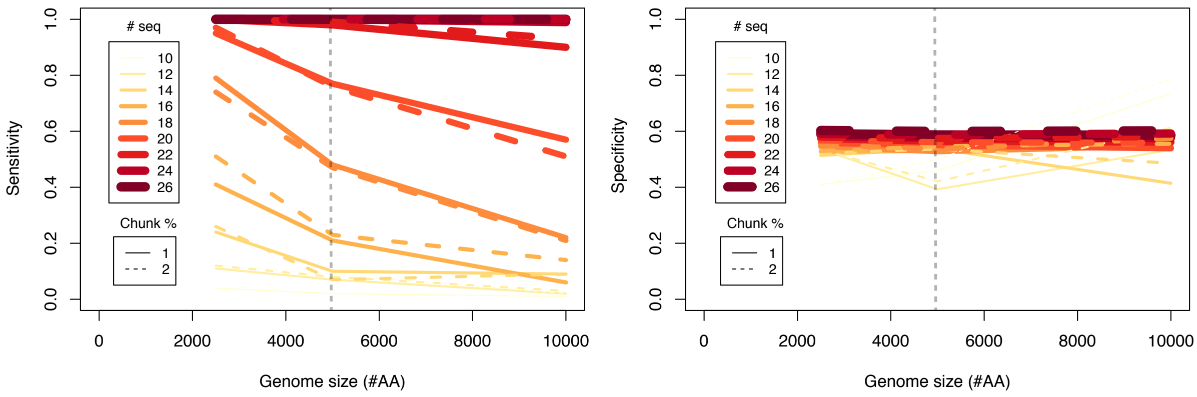


**Fig.S14.** Sensitivity and specificity of the RRF algorithm for the influenza data. Mimicking these data, simulations were performed with no class imbalance, small chunk sizes (either 1 or 2% of the total alignment length), alignment lengths varied between 2500 and 10,000 residues, and including 10 to 26 sequences. The gray vertical line represents the length of the influenza alignment used in this study.

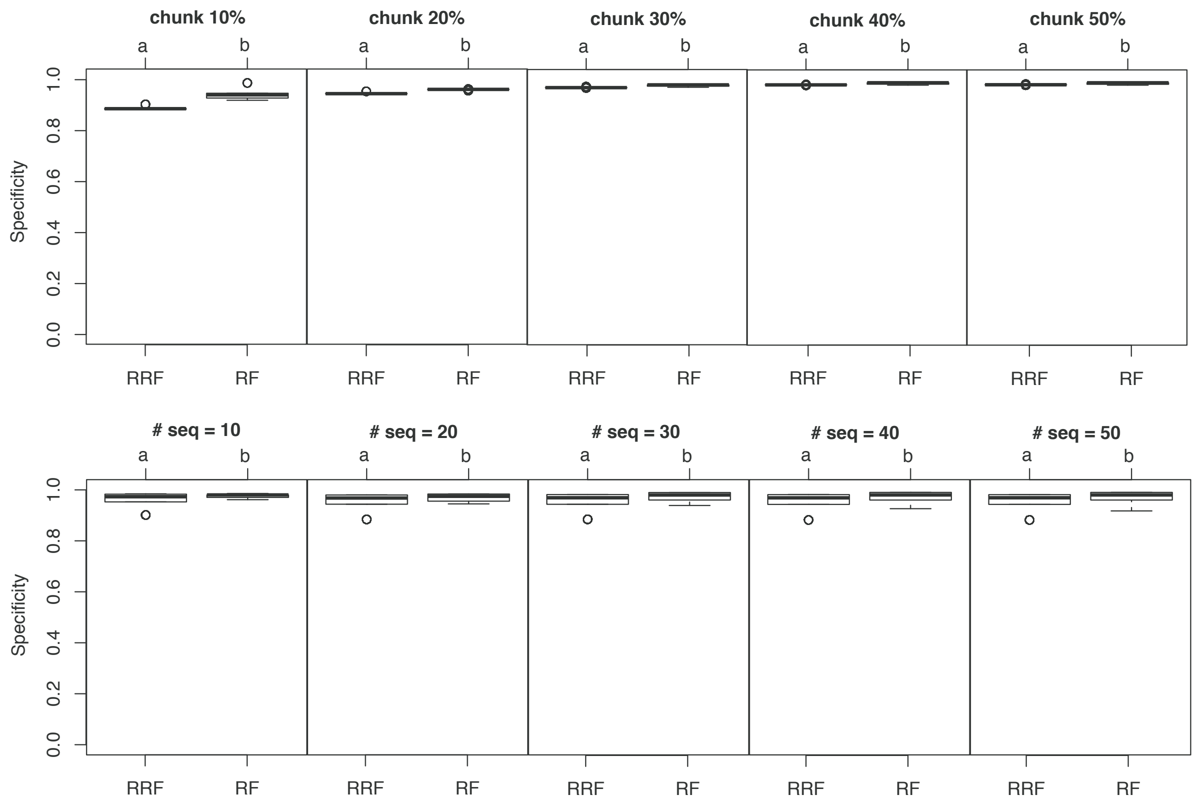


**Fig.S15.** Post-hoc comparisons of specificity between RRF and RF. Boxplots show specificity compared between the RRF and RF algorithms. Results are based on Tukey's Honest Significant Differences, and are derived on an Analysis of Variance fitting specificity as a function the algorithm times number of sequences at different chunk sizes (top row), or chunk size at different number of sequences (bottom row). Results are shown for chunk sizes between 10 and 50%, and for alignments comprising between 10 and 50 sequences. Letter codes indicate significant differences (at the 1% level), with 'a' < 'b.'

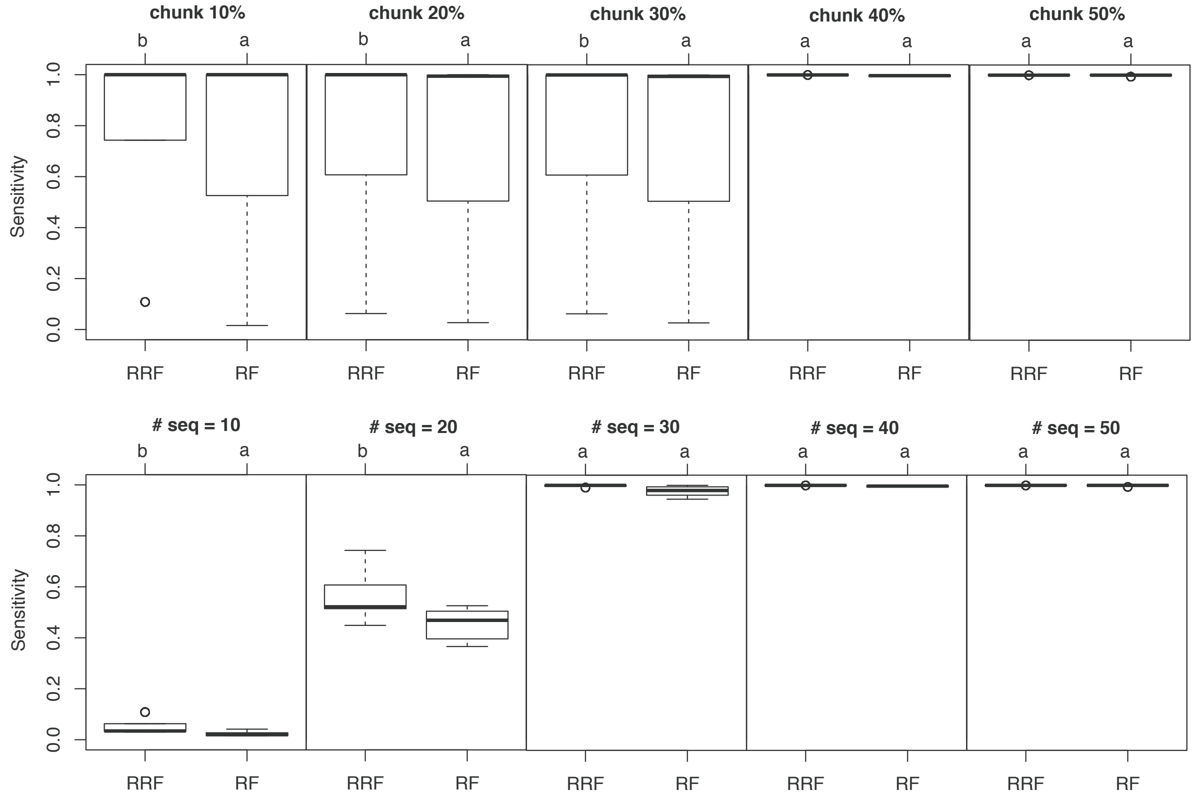


**Fig.S16.** Post-hoc comparisons of sensitivity between RRF and RF. Boxplots show sensitivity compared between the RRF and RF algorithms. Results are based on Tukey's Honest Significant Differences, and are derived on an Analysis of Variance fitting sensitivity as a function the algorithm times number of sequences at different chunk sizes (top row), or chunk size at different number of sequences (bottom row). Results are shown for chunk sizes between 10 and 50%, and for alignments comprising between 10 and 50 sequences. Letter codes indicate significant differences (at the 1% level), with 'a' $<$ 'b.'

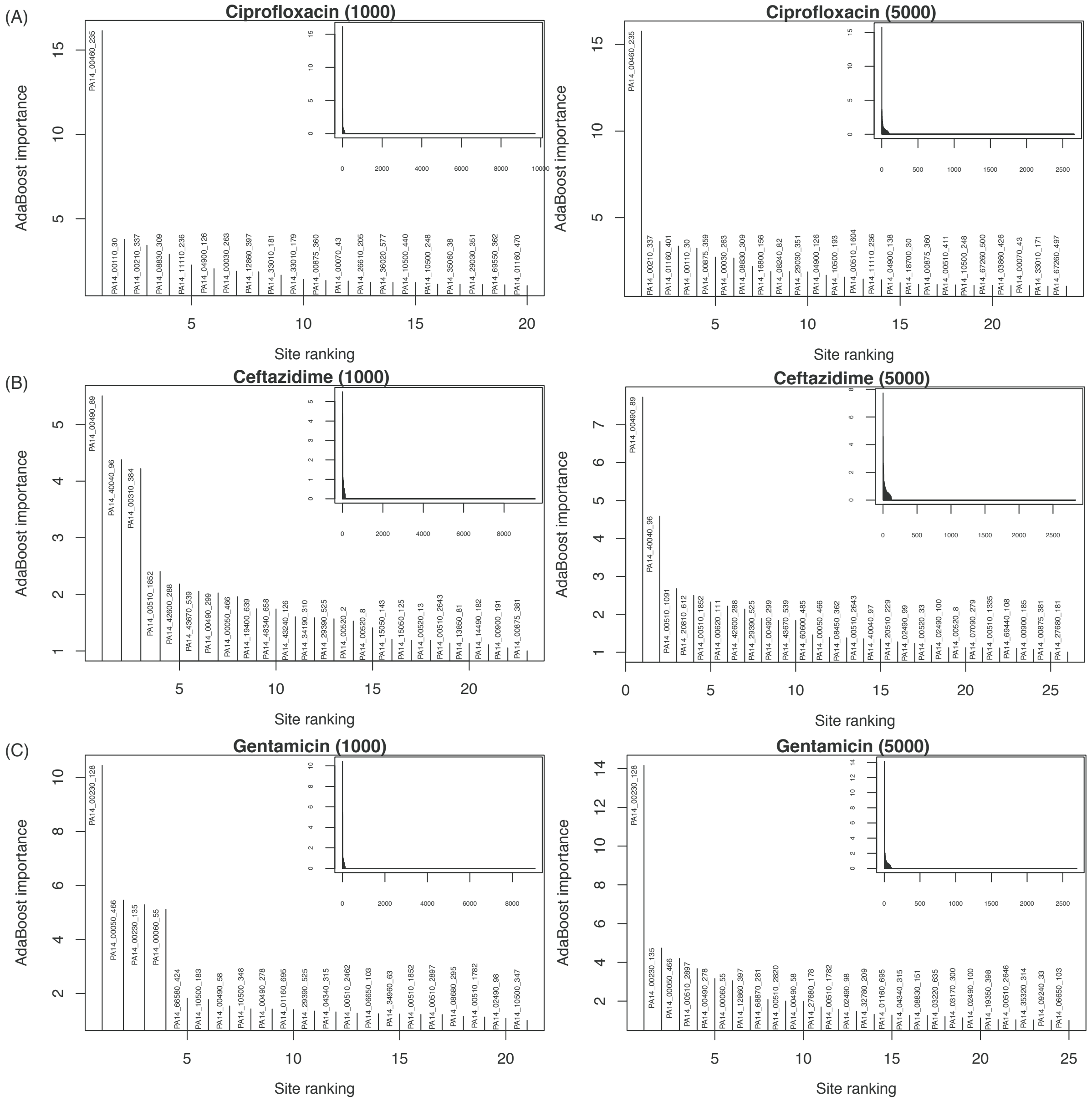


**Fig.S17.** Importance values for the genetic determinants of drug resistance in *P. aeruginosa*. The genes and sites identified are shown for: (A) Ciprofloxacin, (B) Ceftazidime, and (C) Gentamicin. In each panel, the main figure shows the sorted importance values for the most important sites (importance > 1). Results for chunk size of 1000 are shown on the left, and chunk size 5000 (as in Figure 19) on the right. The complete distribution of importance values for all the amino acid positions of interest is shown as insets.

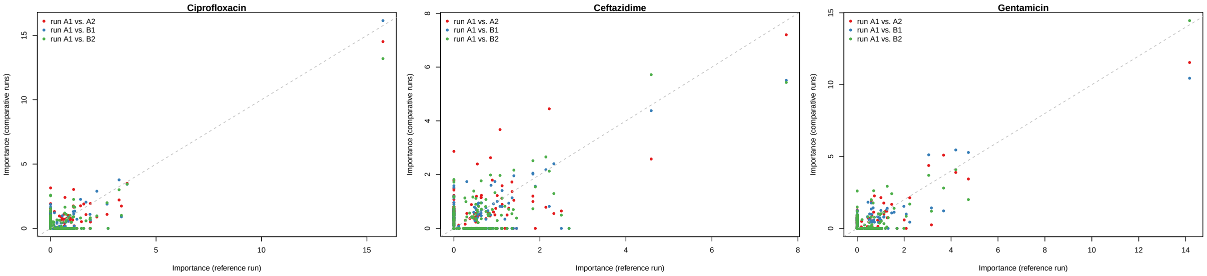


**Fig.S18.** Stability of importance values for the genetic determinants of drug resistance in *P. aeruginosa*. The sites identified are shown for: (A) Ciprofloxacin, (B) Ceftazidime, and (C) Gentamicin. In each panel, the comparison of importance values is between run A1 (replicate 1 under chunk size of 5000, the reference run) and comparative runs A2 (red), B1 (replicate 1 under chunk size of 1000; blue), and B2 (green).

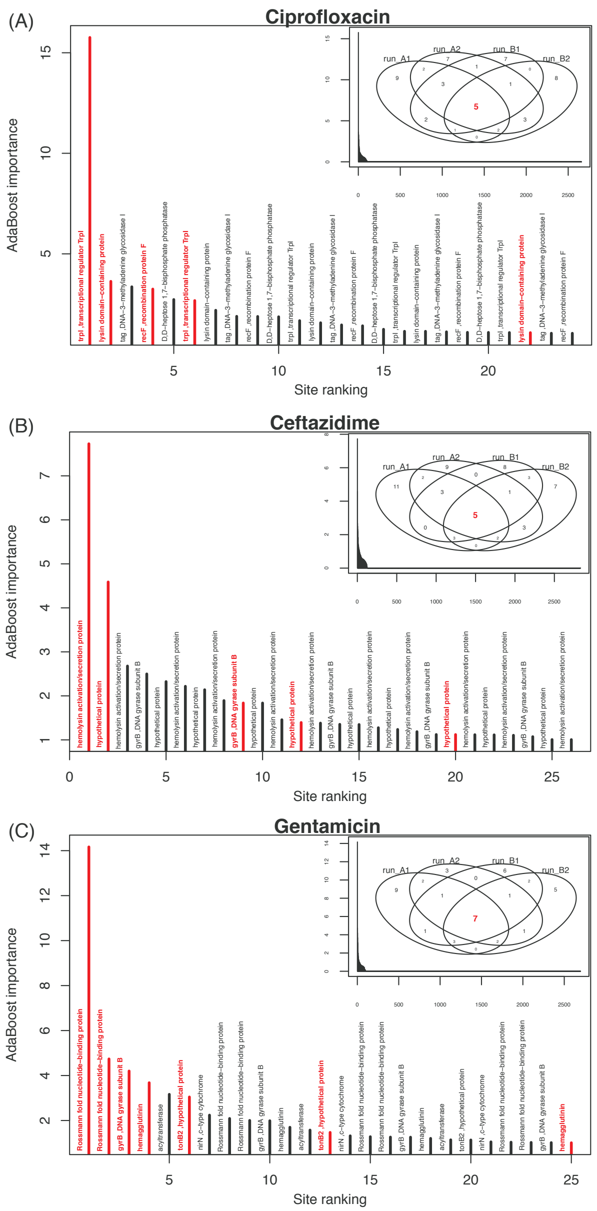


**Fig.S19.** Importance values for the genetic determinants of drug resistance in *P. aeruginosa*. The genes identified are shown for: (A) Ciprofloxacin, (B) Ceftazidime, and (C) Gentamicin. In each panel, the main figure shows the sorted importance values for the most important sites (importance $> 1$) as found in run A1 (replicate 1 under chunk size of 5000). The complete distribution of importance values for all amino acid positions of interest is shown as an inset, which also shows the Venn diagram (intersect) for the top 25 sites found in run A2 (second replicate under chunk size of 5000), and also B1 and B2 (two replicates under chunk size of 1000). The number of sites commonly identified by all four runs is shown in red; the genes in which these sites occur are labeled in red in the main figure. See Figure S17 for details.

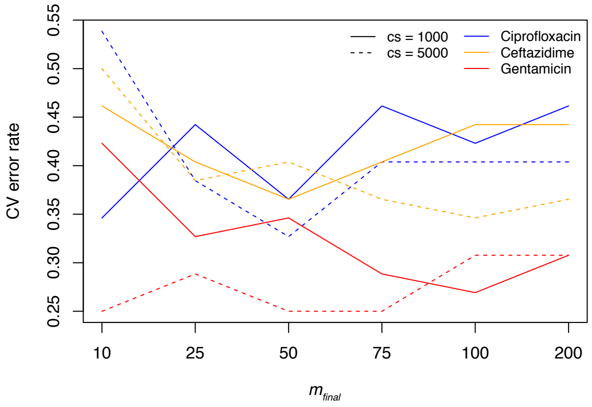


**Fig.S20.** Cross validation errors rates as a function of total number of iteration for the *P. aeruginosa* data. In each case, the mean ten-fold cross-validation error rates were computed over two replicates, and are shown for Ciprofloxacin (blue), Ceftazidime (orange), and Gentamicin (red), with a chunk size of 1000 (solid lines) or 5000 (broken lines).

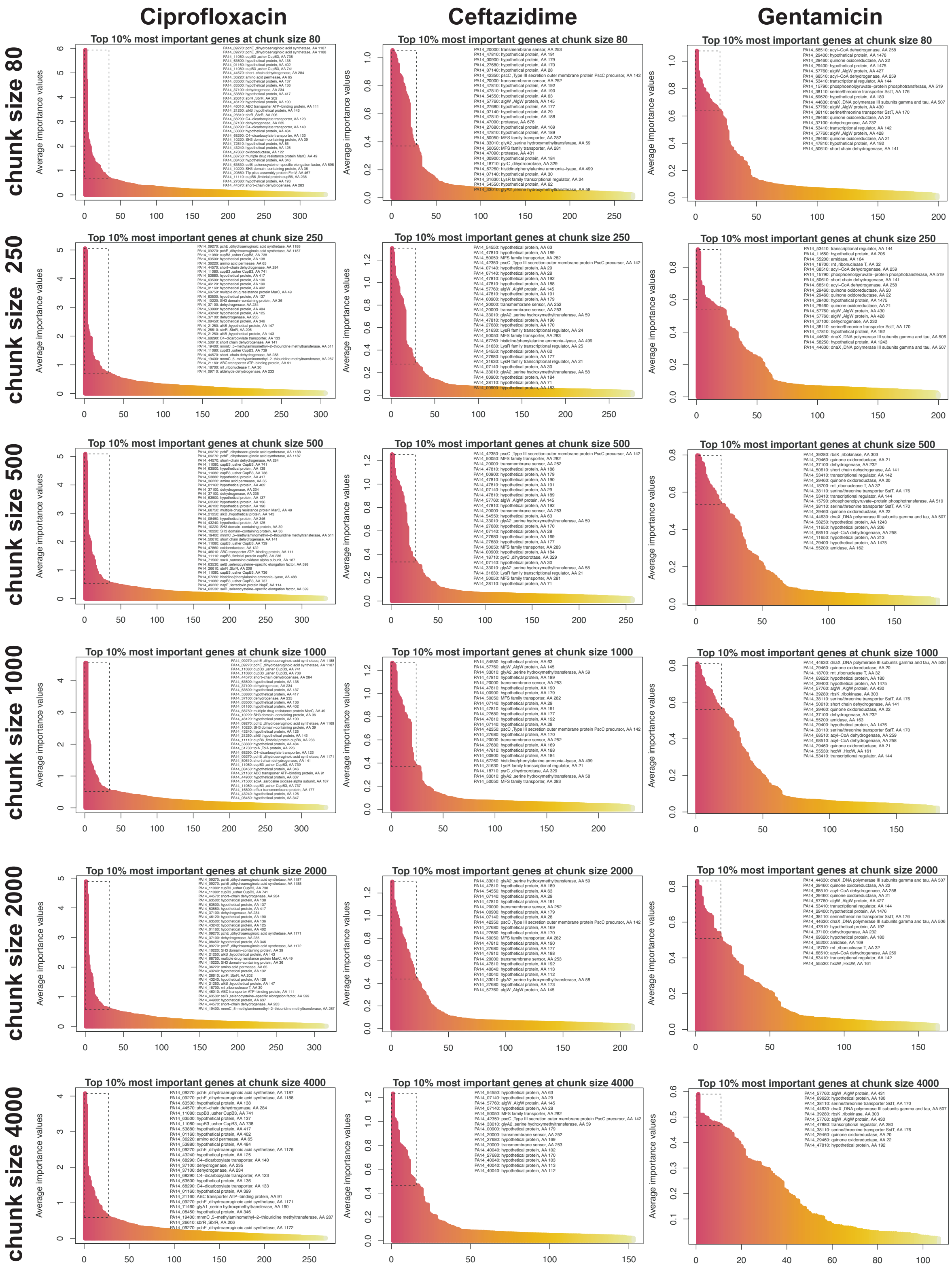


**Fig.S21.** Importance values for the genetic determinants of drug resistance in *P. aeruginosa*. The genes and sites identified are shown for Ciprofloxacin, Ceftazidime, and for Gentamicin. See Figure 5 in main text for details.

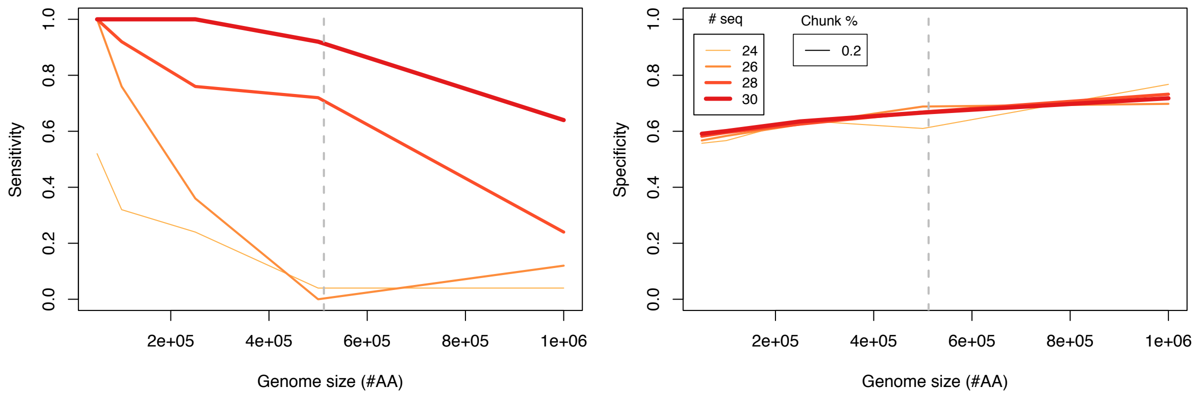


**Fig.S22.** Sensitivity and specificity of the RRF algorithm for the pseudomonas data. Mimicking these data, simulations were performed with no class imbalance, a small chunk size (0.2% of the total alignment length), alignment lengths varied between 50,000 and $10^6$ residues, and including 24 to 30 sequences. The gray vertical line represents the length of the pseudomonas alignment used in this study.
